# Selective Targeting of Pathogenic Tau Seeds via a Novel VHH

**DOI:** 10.1101/2025.01.13.632833

**Authors:** Ankit Gupta, Richard Liu, Devin Keely, Victoria Sunderman, Yogesh Tak, Katerina Kostantoulea, Sandi Jo Terpack, Charles L. White, William P. Russ, Josep Rizo, Nikolaos Louros, Jaime Vaquer-Alicea, Marc I. Diamond

**Affiliations:** Center for Alzheimer’s and Neurodegenerative Diseases, University of Texas Southwestern Medical Center, Dallas, TX 75390; Peter O’Donnell Jr. Brain Institute, University of Texas Southwestern Medical Center, Dallas, TX 75390; Department of Biophysics, University of Texas Southwestern Medical Center, Dallas, TX 75390

**Author notes:** Co-Corresponding Authors: Jaime Vaquer-Alicea, Ph.D. NS8.332, University of Texas Southwestern Medical Center 6124 Harry Hines Blvd. Dallas, TX 75390; Marc I. Diamond, M.D., NS8.334, University of Texas Southwestern Medical Center 6124 Harry Hines Blvd., Dallas, TX 75390. Authors contributed equally to the work (Devin Keely, Victoria Sunderman).

## Abstract

In Alzheimer’s disease (AD) and related tauopathies, progressive pathology has been linked to prion mechanisms, whereby ordered tau assemblies, or “seeds,” form in one cell and transit to neighboring or connected cells where they serve as templates for their own replication. Despite intensive efforts to develop early diagnosis and effective treatment, none has emerged. This is due partly to the structural heterogeneity of tau seeds and limitations in selectively targeting their pathogenic conformations. Here we report the discovery of a camelid variable heavy domain of heavy chain (VHH) that preferentially binds tau seeds of AD, corticobasal degeneration (CBD), and PS19 tauopathy mouse brains. From a published synthetic VHH yeast display library, we identified VHH clones that bound 2N4R tau monomer. After counter-screening for immunoprecipitation of seeding from human brain, we identified two seed-selective anti-tau VHH—VHH(510) and VHH(50)—that preferentially bound pathological tau. We enhanced the stability of these VHHs through framework mutations without affecting their seed-binding characteristics. We characterized VHH(510) in detail, determining that it binds the carboxy terminus of tau, maintains robust seed avidity even in the presence of competing monomer, and stains pathological tau inclusions in mouse and human AD tissues. These results highlight the power of seed-selective VHH to bind pathogenic tau, paving the way for future therapeutic and diagnostic applications across a wide range of neurodegenerative disorders.

## Introduction

Tau aggregation in amyloid fibrils causes neuronal dysfunction, cell death, and cognitive decline. This pathology underlies a spectrum of neurodegenerative disorders collectively referred to as tauopathies, including Alzheimer’s disease (AD), progressive supranuclear palsy (PSP), corticobasal degeneration (CBD), and Pick’s disease (PiD)(*1*). Each disease typically exhibits a unique conformation of assembled tau (*2*, *3*). We originally proposed that distinct fibrillar aggregates, or strains, function as prions, promoting progressive pathology by serving as templates that corrupt native tau (*4–6*). The propagation of tau seeds between neurons is considered a key driver of disease progression (*7*, *8*). Consequently, targeting tau seeds represents a promising strategy for early disease detection and intervention.

Various therapeutic approaches have been investigated over the last decade to inhibit tau pathology (*9–11*). While multiple antibodies have been developed to target different regions or modifications of tau—including the N-terminal region, proline-rich domain, repeat domain, and phosphorylated or acetylated epitopes (*12–16*)— none have yet demonstrated efficacy in patients. Failures likely stem from the heterogeneity of tauopathies, off-target effects, barriers to the efficient delivery of therapeutics to the brain, and the challenge of targeting relevant pathogenic tau conformations.

In this context, camelid variable domain of heavy chain (VHH) peptides have gained attention for their small size, stability, straightforward production, and specificity (*17*, *18*). These features make them excellent reagents to bind difficult targets such as G-protein-coupled receptors (GPCRs), enzymes, amyloids, and viral proteins. Their ability to penetrate tissues and potentially cross the blood-brain barrier further underscores their utility for targeting neurodegenerative diseases (*19–24*). However, many anti-tau VHHs generated to date target recombinant tau monomer or heparin-induced fibrils, which can differ structurally from aggregates observed in the disease (*2*, *25*). Thus, reagents that preferentially bind human-brain-derived tau seeds may offer greater therapeutic promise, as each tauopathy exhibits its own structural “fingerprint” (*3*, *26*).

Despite notable progress, our understanding of the biochemical and structural properties of tau seeds remains limited, and isolating large quantities of highly purified patient-derived tau seeds is impractical for large screening campaigns. To address this challenge, we devised a two-stage “hybrid” VHH screening strategy. We first enriched VHHs that bind full-length 2N4R tau monomer from a synthetic yeast display library and then employed immunoprecipitation to identify clones that preferentially bound tauopathy brain-derived tau seeds. This enabled us to discover anti-tau VHHs that do not strongly bind heparin-induced recombinant fibrils or tau monomer but do bind disease-derived tau seeds. These VHHs retained seed binding even in the presence of excess monomer. One, VHH(510) mapped to the C-terminus of tau, immunoprecipitating AD tau seeds, discriminating disease and control brain lysates, and staining tau inclusions in both a P301S mouse model and human AD tissue. This approach has led to the generation of seed-selective VHHs with potential utility for research, diagnostic, and therapeutic applications in AD and other tauopathies.

## Results

### Two-Step VHH Screening Identifies Tau Seed–Binding Clones

Our primary goal was to identify VHHs that preferentially recognize pathogenic tau seeds from AD brains. Given the challenges of obtaining large quantities of pure tau seeds from human sources, we devised a hybrid screening strategy to generate VHHs that bind tau monomer while also recognizing authentic tau seeds (Figure 1). In the first step, we enriched VHH clones from a fully synthetic yeast display library (∼1×10^8^ clones; (*27*)) that bound full-length 2N4R tau monomer. We performed five rounds of selection—two rounds of Magnetic-Activated Cell Sorting (MACS) followed by three rounds of Fluorescence-Activated Cell Sorting (FACS)— progressively lowering the tau concentration from 500 nM to 10 nM to isolate high-affinity clones (Figure 1A, Figure S1). Following these selections, we obtained ∼900 yeast clones with verified tau monomer binding.

**Figure 1.**
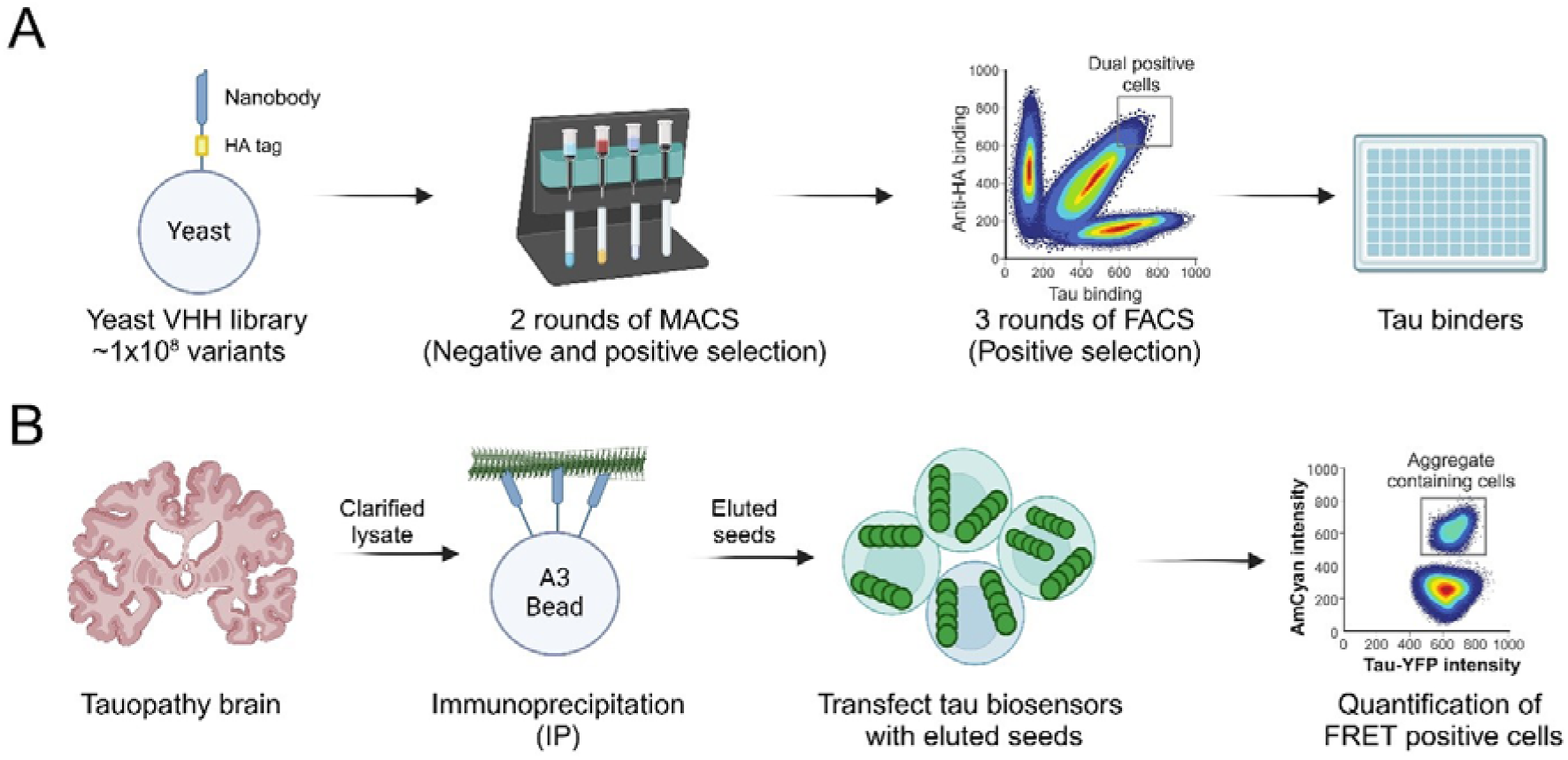
A hybrid approach for VHH screening. (**A**) Schematic of the initial screen using a yeast surface-display library of approximately 1×10^8^ VHH clones (procured from Kerafast, Cat EF0014-FP). Two rounds of magnetic-activated cell sorting (MACS) enriched for yeast displaying VHHs that bind full-length 2N4R tau, followed by three rounds of fluorescence-activated cell sorting (FACS) to further refine the selection. See **Figure S1** for additional details on screening parameters. (**B**) Strategy to identify VHHs that preferentially recognize seed-competent tau. VHHs identified from (**A**) were used to immunoprecipitate tau seeds from clarified AD brain lysates, and the eluted material was transfected into tau biosensor cells. FRET-positive cells in this assay reflect pathogenic seeding activity, thereby serving as a readout of seed binders.

In the second step, we screened these clones for seed-binding by immunoprecipitation (IP) from patient-derived lysates (Figure 1B). We used clarified homogenates of the frontal cortex from two AD cases (AD1, AD2), a CBD case, and a P301S mouse model, each known to harbor distinct pathogenic tau seeds. After incubation, we eluted bound material and measured seeding activity in v2L tau biosensor cells (*28*, *29*). This approach allowed us to prioritize clones capable of immunoprecipitating pathologically relevant tau conformations. From ∼900 starting clones, ∼300 robustly pulled-down tau seeds (Figure S2). Sequence analysis of the best clones revealed >15 unique VHHs, including VHH(510) and VHH(50). We selected these highly enriched clones for further characterization.

### Selected VHHs Preferentially Bind AD and CBD-Derived Tau Seeds

To assess whether these VHHs were selective for human brain-derived seeds, we performed a more detailed screen using IP-and-seeding assays against multiple tau sources (Figure 2A).

**Figure 2.**
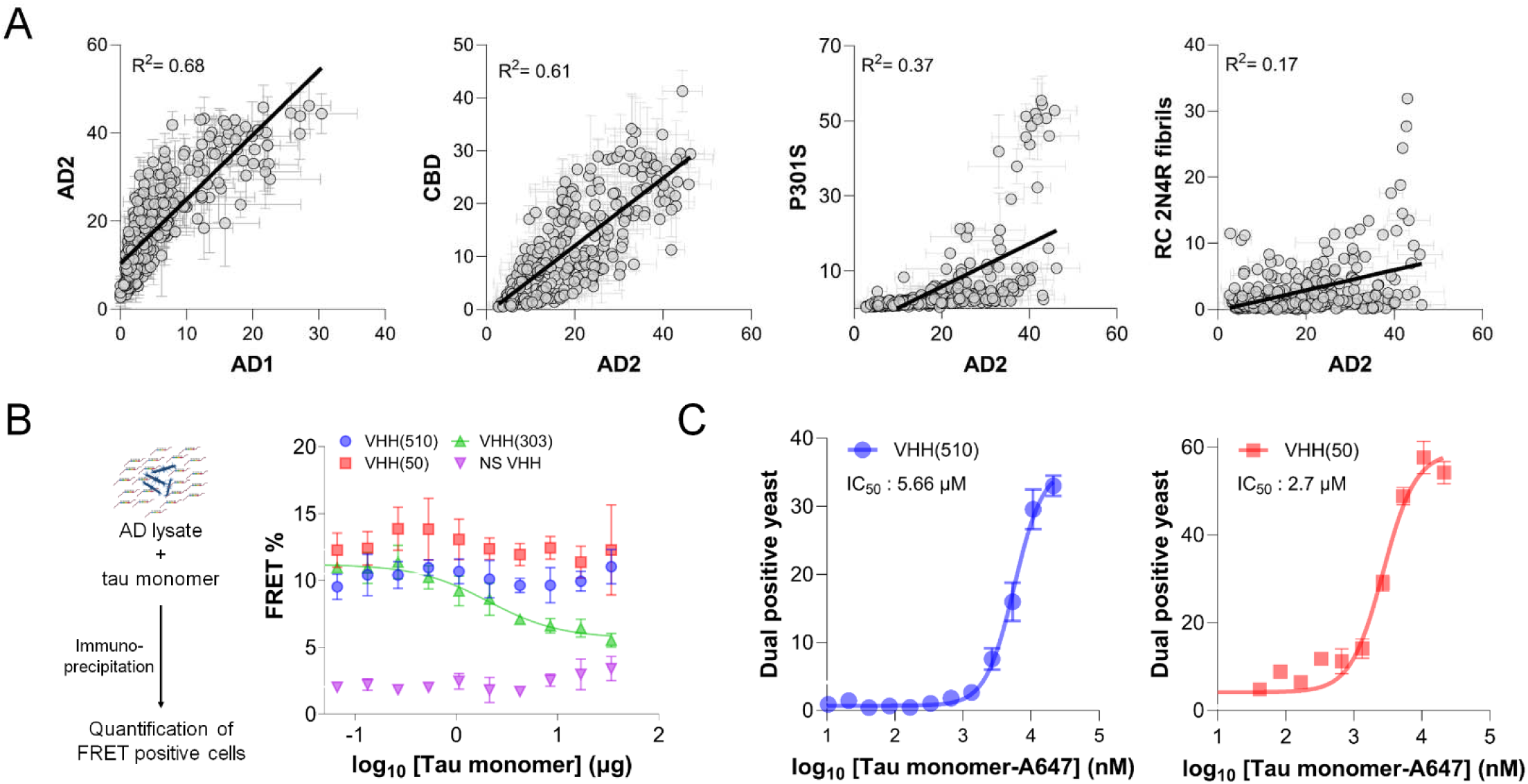
Discovery of VHHs selective for AD seeds. (**A**) Correlation between the percentage of FRET-positive cells detected by an IP-and-seeding assay for the top 300 VHH clones, tested with distinct tau seed sources. Shown are pairwise comparisons between two independent AD samples (leftmost panel), an AD and a CBD sample (second panel), an AD sample and P301S lysate (third panel), and an AD sample with heparin-induced recombinant fibrils (rightmost panel). **Figures S2** and **S3** provide additional data and analyses. (**B**) Left: schematic of a competition experiment testing whether excess monomeric 2N4R tau interferes with VHH-mediated pull-down of AD-derived seeds in an IP-and-seeding assay. Right: the resulting FRET signal observed in v2L tau biosensor cells for anti-tau VHH(510), VHH(50), and VHH(303), as well as a non-specific VHH (NS VHH), across increasing concentrations of monomeric tau. (**C**) Determination of IC_50_ values for anti-tau VHH(510) (left) and VHH(50) (right) against Alexa-647–labeled 2N4R monomer. Here, the percentage of dual-positive yeast cells (i.e., cells displaying a given VHH on their surface and binding the labeled tau) is plotted against increasing concentrations of fluorophore-labeled tau.

Specifically, we compared two independent AD cases, one CBD case, P301S mouse brain lysate, and heparin-induced recombinant 2N4R and K18 fibrils. We quantified seeding as the percentage of FRET-positive cells in the v2L tau biosensor line (*29*). Scatter plots (Figure 2A; Figure S3) revealed strong correlations in seed binding among AD and CBD samples but weak or negligible correlations between patient-derived tau seeds and either P301S or recombinant fibrils (Figure S3D-I). These findings reinforced previous structural evidence that fibrils derived from human tauopathy brains differ significantly from heparin-induced recombinant fibrils (*2*, *25*) Seed Binding is Maintained in the Presence of Excess Monomer

A critical criterion for any tau-directed therapeutic reagent is to recognize pathogenic seeds selectively, rather than the abundant soluble tau monomer found throughout the human body. We therefore performed a competition experiment in which anti-tau VHHs were incubated with an excess (5x) of 2N4R monomer during IP from AD brain lysates (Figure 2B). The pulled-down material was then eluted and assayed in tau biosensor cells. Both anti-tau VHH(510) and VHH(50) retained their robust seeding signal, indicating that excess monomer did not compete away pathological seeds. In contrast, known monomer-selective VHH(303) exhibited markedly lower seed binding under these conditions. These data indicated that VHH(510) and VHH(50) preferentially bound pathological tau seeds.

Next, we carried out on-yeast IC_50_ measurements to quantify monomer affinity (Figure 2C). Both VHH(510) and VHH(50) had micromolar-range affinities for tau monomer, with VHH(510) notably weaker (higher IC_50_). Together, these results prompted us to focus on VHH(510) for more detailed investigations.

### Framework Optimization Improves VHH Expression and Stability

To produce larger quantities of these VHHs for functional and structural assays, we cloned their coding sequences into *E. coli* expression vectors. However, the wild-type variants were challenging to purify as they tended to aggregate at high concentrations during expression and purification (Figure S4 and S5), limiting yields of soluble protein. Following a strategy described previously (*30*), we introduced framework mutations that bolstered VHH stability without altering the antigen-binding complementarity-determining regions (CDRs). These stabilized mutant variants are referred to as VHH(50M) and VHH(510M). SDS-PAGE analyses confirmed that VHH(50M) and VHH(510M) expressed at higher levels and exhibited diminished aggregation (Figure S4 and S5). After purification via Amsphere™ A3 beads (Figure S6), we measured secondary structure of VHH by circular dichroism (CD). Both stabilized VHHs retained a typical β-sheet signature and were stable until ∼55 °C (Figure S7). When tested by IP-and-seeding (Figure 3), the framework-optimized VHHs preserved the seed-binding capabilities of their wild-type counterparts, validating that the mutations did not compromise specificity.

**Figure 3.**
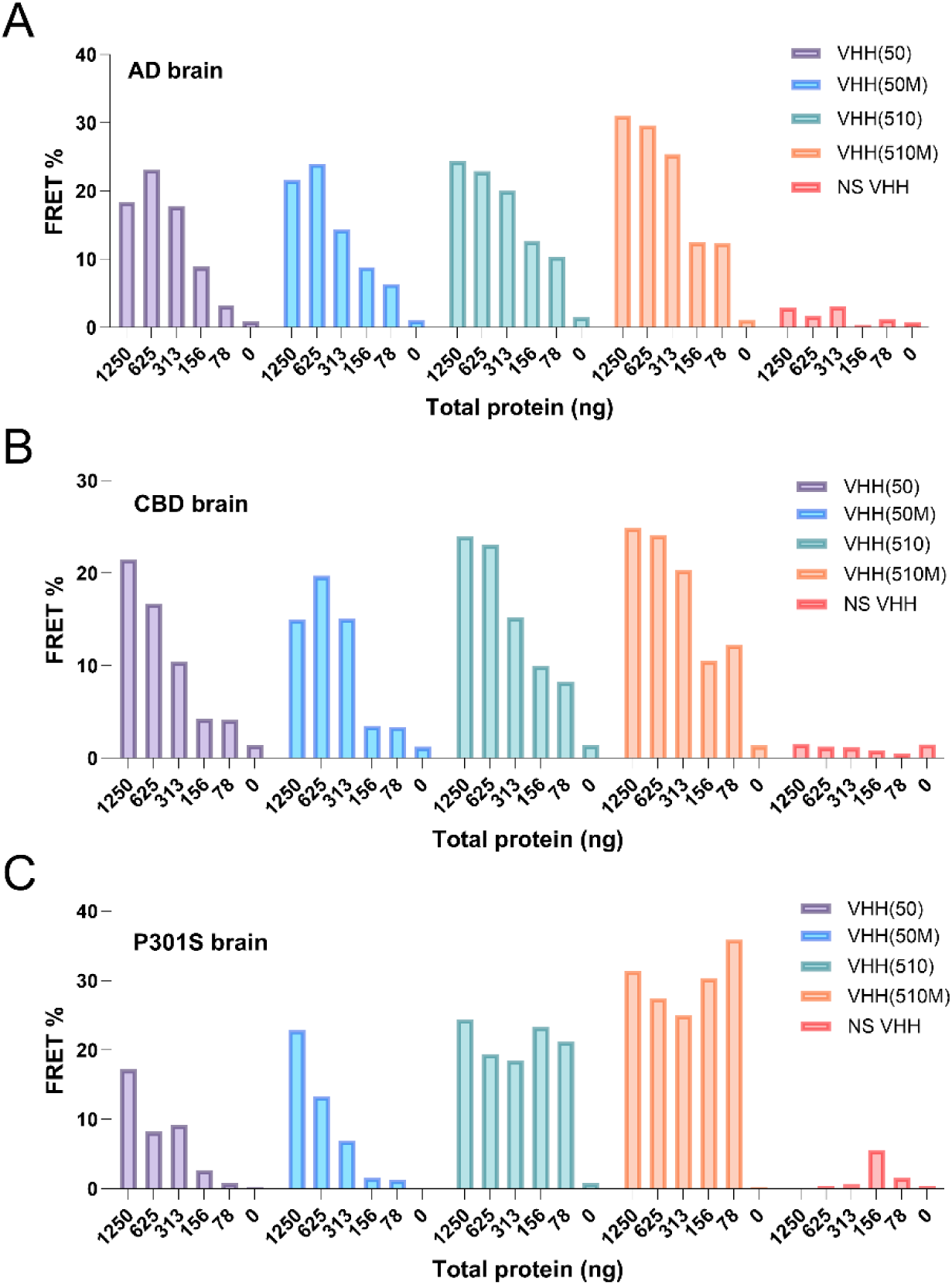
Comparison of seed-binding by wild-type (WT) and stabilized (M) anti-tau VHHs. (**A**-**C**) Immunoprecipitation was performed with purified wild-type VHHs (anti-tau VHH(50) and VHH(510)) and their stabilized counterparts (anti-tau VHH(50M) and VHH(510M)) using Amsphere™ A3 resin across different concentrations of lysate total protein. Bar graphs show the resulting FRET% from v2L tau biosensor cells following IP and elution from (**A**) AD brain lysate, (**B**) CBD brain lysate, and (**C**) P301S mouse brain lysate. A non-specific (NS) VHH was included as a negative control.

### Anti-Tau VHH(510M) Selectively Immunoprecipitates Seeds from AD and CBD brains

We further characterized the binding properties of the anti-tau VHHs VHH(510M) and VHH(50M) by performing immunoprecipitation (IP) with purified VHHs against brain samples from multiple cases of Alzheimer’s disease (AD), a Corticobasal degeneration (CBD) case, and from a P301S mouse (Figure 4). Here, soluble, clarified homogenates of frozen frontal cortex tissue from several AD and CBD brains were prepared, and anti-tau VHH(510M) and VHH(50M) were used to pull down tau seeds using Amsphere™ A3 beads. We observed that both nanobodies successfully immunoprecipitated tau seeds from AD and CBD brains, and these eluted seeds were detectable on v2L tau biosensor cells (Figure 4A, B). Additionally, both VHHs bound to P301S brain samples but showed no binding to control brain samples (Figure 4C, D). Interestingly, the seed binding was comparable for both VHH(510M) and VHH(50M), with VHH(510M) demonstrating slightly superior binding compared to VHH(50M).

**Figure 4.**
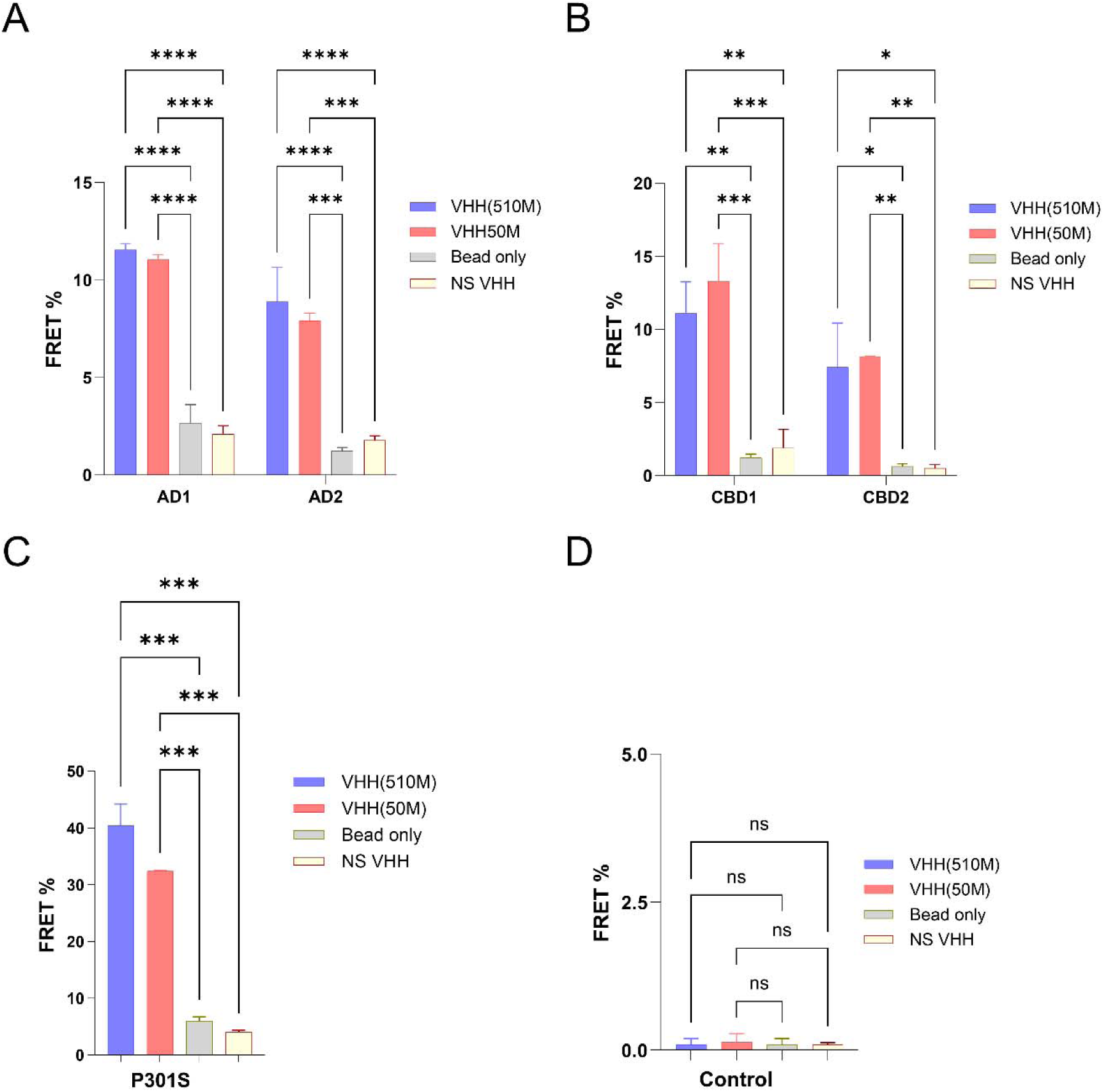
Immunoprecipitation of tau seeds from various tauopathy and control brains. (**A**-**C**) Seed capture from two AD brain samples (AD1, AD2), two CBD brain samples (CBD1, CBD2), and a P301S mouse brain, respectively, using anti-tau VHH(510M) or VHH(50M). (**D**) Immunoprecipitation from a non-tauopathy control brain. Bars indicate mean ± S.D. (n=3). Significance was evaluated against a non-specific VHH (NS VHH) and bead-only controls using two-way ANOVA with Tukey’s multiple comparison test (AD, CBD) or one-way ANOVA with Šídák’s multiple comparisons test (P301S). ***p ≤ 0.0002; **p ≤ 0.001; ****p ≤ 0.0001.

To further validate the observation of preferential seed binding, we performed dot-blot analysis and assessed the binding of VHH(510M) and VHH(50M) to AD, P301S, and control brain samples (Figure 5). We blotted varying concentrations of brain lysates onto membranes and detected tau using the VHHs. As expected, VHH(510M) showed binding to only tauopathy brains and minimal binding to control brains (Figure 5A, B left panel), whereas VHH(50M) bound to both AD and control brains (Figure 5A, B right panel). These observations align well with our affinity measurements, where VHH(510M) shows ∼8-fold weaker binding to tau monomer than VHH(50M). Overall, these results suggest that VHH(510M) has a high affinity for tau seeds and a lower affinity for tau monomers, while VHH(50M) exhibits a comparatively higher affinity for the monomeric form of tau.

**Figure 5.**
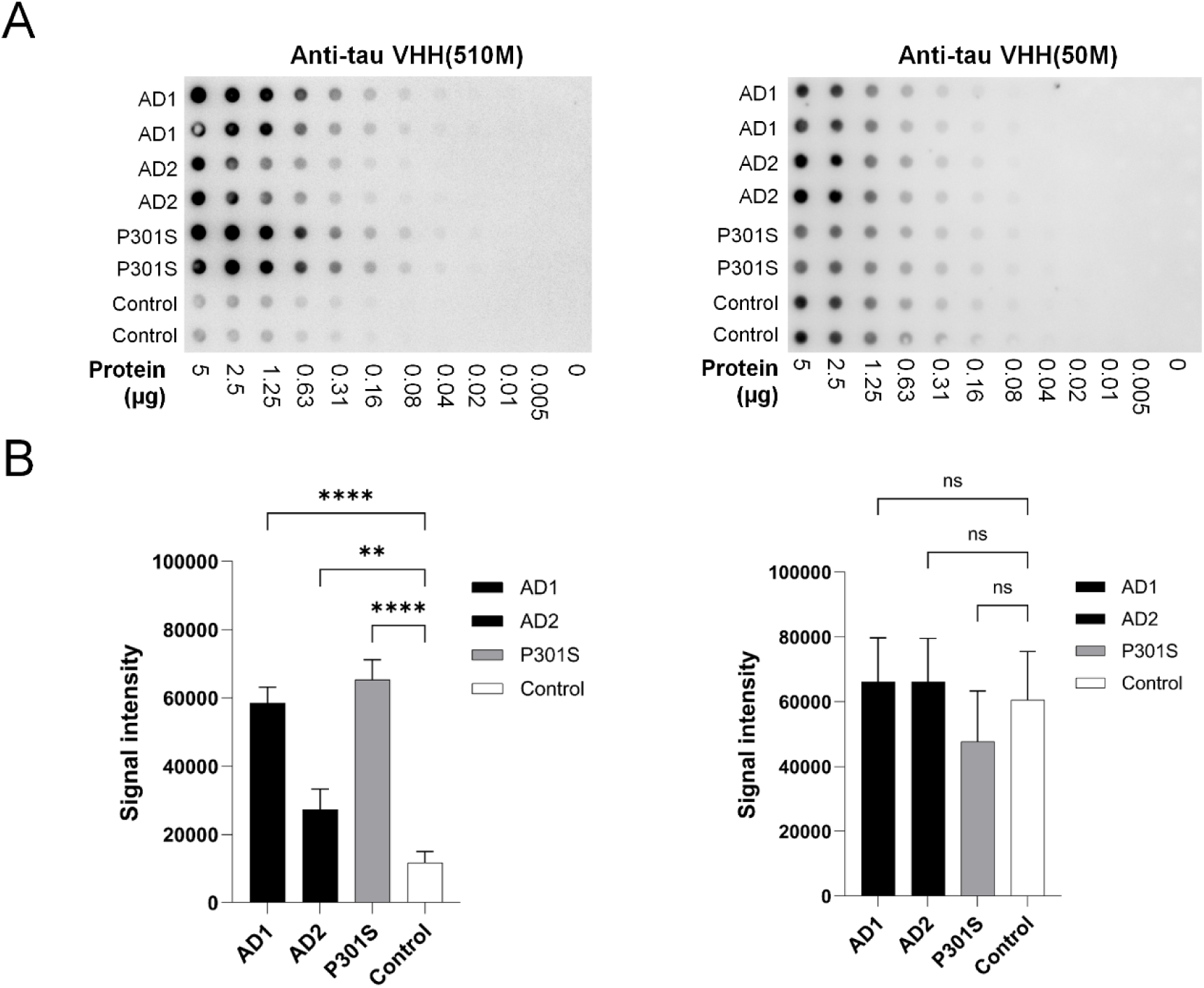
Dot blot analysis of anti-tau VHH binding to AD, P301S, and control brain lysates. (**A**) Dot blots using anti-tau VHH(510M) (left) and anti-tau VHH50M (right) as detection molecules against serial dilutions of lysates from two brains of AD cases (AD1, AD2), a P301S mouse model, and control. (**B**) Quantification of signal intensity at 1.25 µg of total protein, based on integrated optical densities measured with Fiji (version 1.54f). Anti-tau VHH(510M) (left panel) preferentially binds to AD and P301S lysates while showing minimal reactivity to control tissue. By contrast, VHH50M (right panel) shows broadly similar signal across all tested samples. Error bars represent S.D. (n = 4). Statistical significance was assessed with one-way ANOVA followed by Šídák’s multiple comparisons test (p ≤ 0.0029, **p ≤ 0.0001).

### Anti-Tau VHH(510M) has Low Affinity for Monomeric Tau

To confirm that the weak monomer affinity observed on yeast carried through to the purified VHHs, we used Flow-Induced Dispersion Analysis (FIDA) (Figure 6B, Figure S8). In these experiments, tau monomer was titrated with a fixed concentration of fluorescently labeled VHH, and their binding affinities were determined. The binding affinities of the purified VHHs calculated from the in-solution kinetics are consistent with our "on-yeast IC_50_" measurements. Our kinetic measurements showed that both VHHs exhibited poor affinity for the tau monomer, with K_D_ values in the micromolar range. Specifically, the VHH(510M) demonstrated a K_D_ of ∼4 µM (Figure 6B, and S8A), that is comparatively higher than what we observed for VHH(50) (K_D_ for VHH(50M): ∼0.6 µM) (Figure S8B). Together these results suggested that the seed-binding anti-tau VHHs had weak affinity for tau monomer.

**Figure 6.**
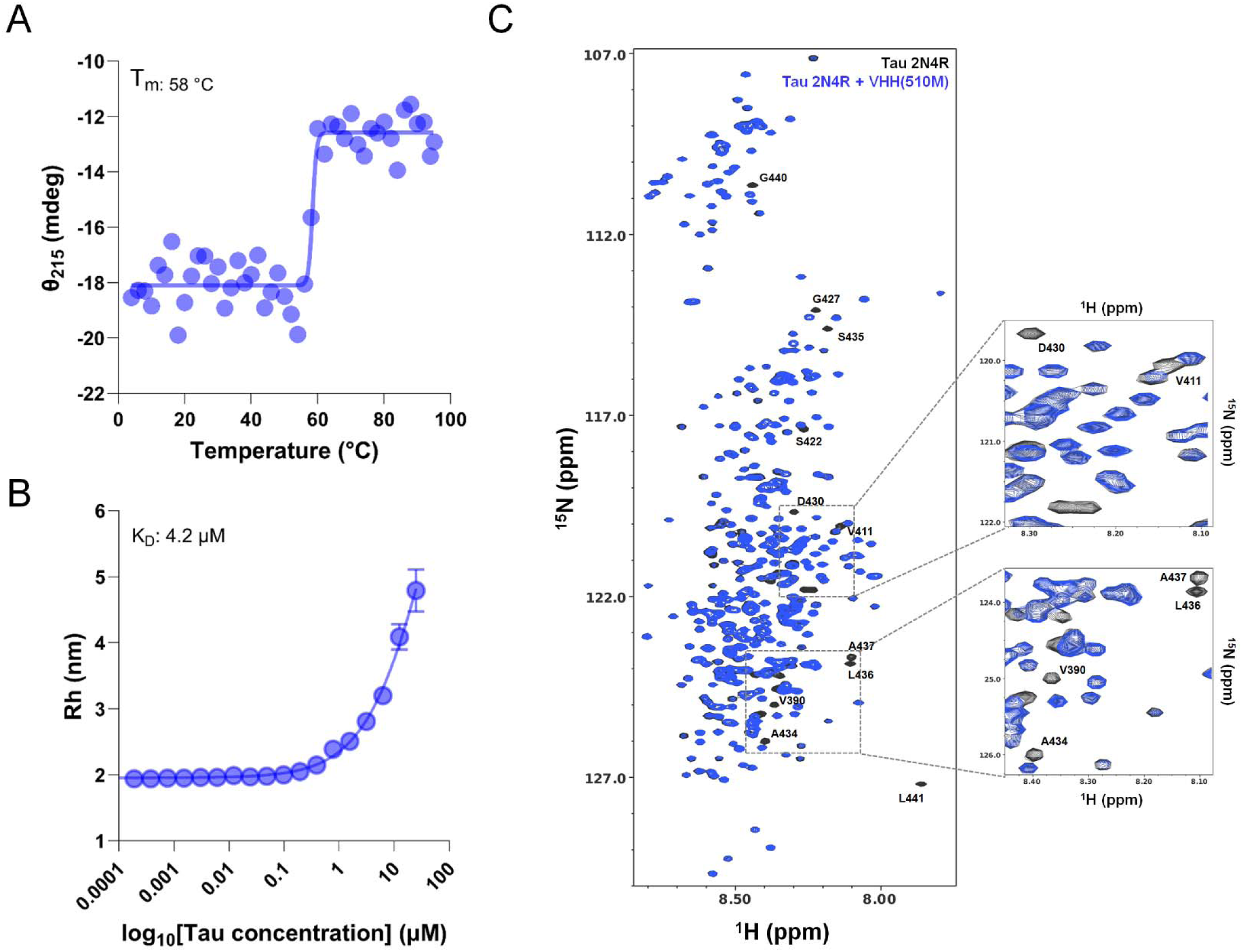
Stability measurements and epitope mapping of anti-tau VHH(510M). (**A**) Representative thermal denaturation profile of VHH(510M) monitored by circular dichroism (CD) at 215 nm (θ_215_), measured from 4 °C to 95 °C. Data were fitted to a sigmoidal model to estimate the midpoint of thermal unfolding (T_m_). See **Figure S7** for additional details. (**B**) In-solution affinity measurement of VHH(510M) using Flow-Induced Dispersion Analysis (FIDA). The hydrodynamic radius (R_h_) of fluorescently labeled VHH(510M) was plotted against increasing concentrations of tau 2N4R monomer. Fitting to a 1:1 binding model yielded a K_D of ∼4.2 µM. See **Figure S8** for more information. (**C**) Overlaid 2D ^1^H,^15^N-HSQC spectrum of ^15^N-labeled tau 2N4R alone (black) and in the presence of equimolar VHH(510M) (blue). Several peaks—particularly in the C-terminal region—shift or disappear upon binding, indicating that (VHH510M) recognizes residues at the carboxy-terminus of tau. Insets show enlarged views of selected regions highlighting peak perturbations.

### Anti-tau VHH(510M) Binds to the Carboxy-Terminus of Tau

We next used solution-state NMR to localize the binding epitope of VHH(510M) on tau. We recorded the ^1^H,^15^N-HSQC spectra of full-length tau 2N4R as described previously (*31*) and overlapped the HSQC peaks with previously published NMR spectra of tau (BMRB Entry 50701) to assign the peaks. Next, we incubated the purified anti-tau VHH(510M) with ^15^N labeled full-length tau 2N4R at 1:1 molar ratio and recorded the ^1^H,^15^N-HSQC spectrum. The superimposed two-dimensional ^1^H,^15^N-HSQC spectra of tau alone and with anti-tau VHH(510M) (Figure 6C) showed broadening or disappearance of several cross-peaks—especially those corresponding to residues near the C-terminus—pinpointing its epitope.

### Anti-tau VHH(510M) Stains Tau Inclusions in P301S Mice and Human AD brains

Finally, we assessed the ability of VHH(510M) to detect intracellular tau aggregates in formalin-fixed tissues from a PS19 P301S tauopathy mouse model (*32*) and from postmortem human AD brains. We stained the brain tissue of PS19 mice at various ages (3 to 12 months) with fluorescently conjugated anti-tau VHH(510M). We also used the AT8 antibody, a well-established antibody that stains phosphorylated tau as a control to compare the staining pattern. We observed that the anti-tau VHH(510M) stained phosphorylated tau inclusions in P301S mice as early as 3 months. VHH(510) produced strong staining in various regions of the brain including the neocortex, amygdala, hippocampus, and brain stem. Interestingly, the staining pattern of anti-VHH(510M) overlaps well with the AT8 antibody as we observe substantial colocalization of both molecules. However, we find that VHH(510M) detects additional inclusions in various brain regions (Figure 7 and S9), suggesting that it may detect certain non-phosphorylated tau species. Notably, these inclusions are abundantly present in the brainstem, as previously described (*33*)

**Figure 7.**
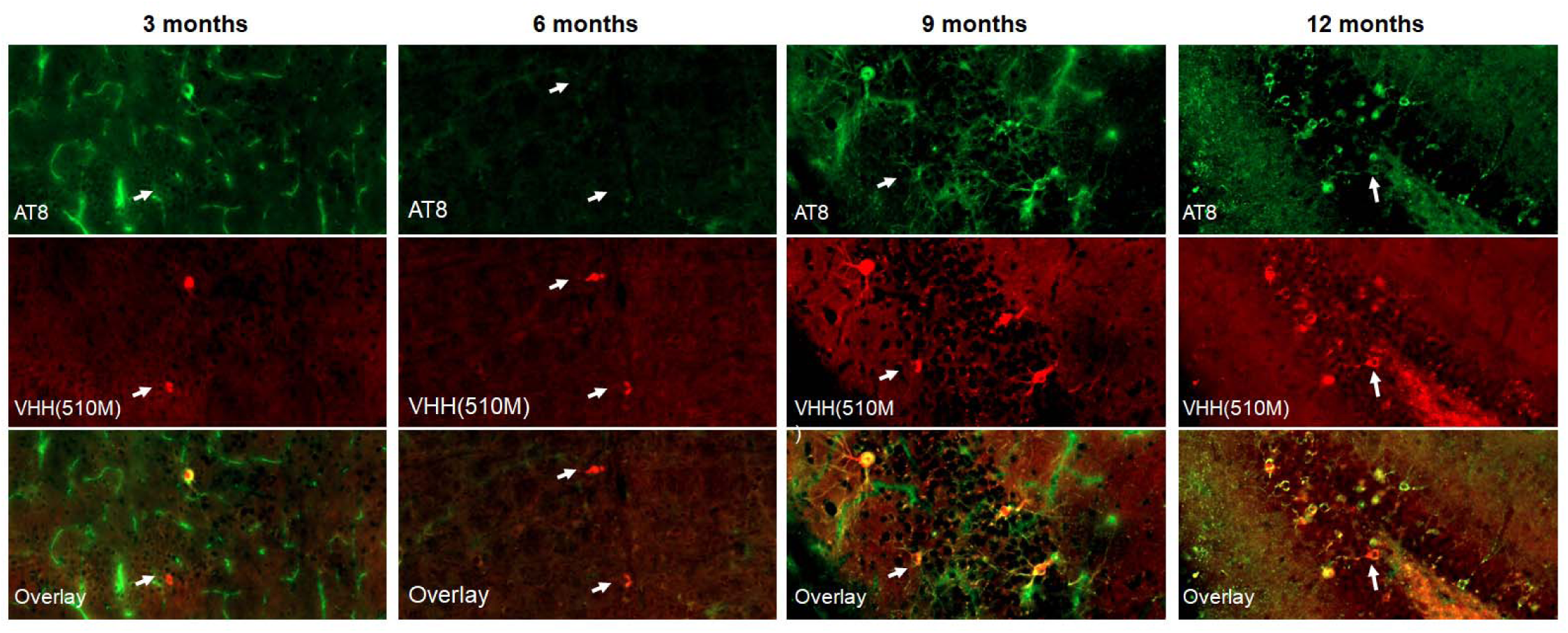
Age-dependent immunostaining of P301S mouse brains with VHH(510M). Shown are representative images from 3-, 6-, 9-, and 12-month-old P301S tauopathy mice stained with the phospho-tau antibody AT8 (green) and anti-tau VHH510M (red). Anti-tau VHH510M labels pathological inclusions that overlap with AT8-positive aggregates but also highlights additional inclusions not recognized by AT8 (white arrows). These observations suggest that VHH(510M) detects both phospho-tau and non-phosphorylated forms of aggregated tau. See **Figure S9** for additional images and details.

In human AD tissue microarray sections (Figure 8), VHH(510M) overlapped with AT8 in many inclusions, including pretangles, mature neurofibrillary tangles, and ghost tangles. However, it showed minimal labeling of neuritic plaques or tangle-associated neuritic clusters, suggesting a selectivity for a subset of pathogenic tau conformations. Together, these findings complement the IP-and-seeding data, confirming that VHH(510M) selectively recognizes pathological tau in a mouse model and human AD tissue.

**Figure 8.**
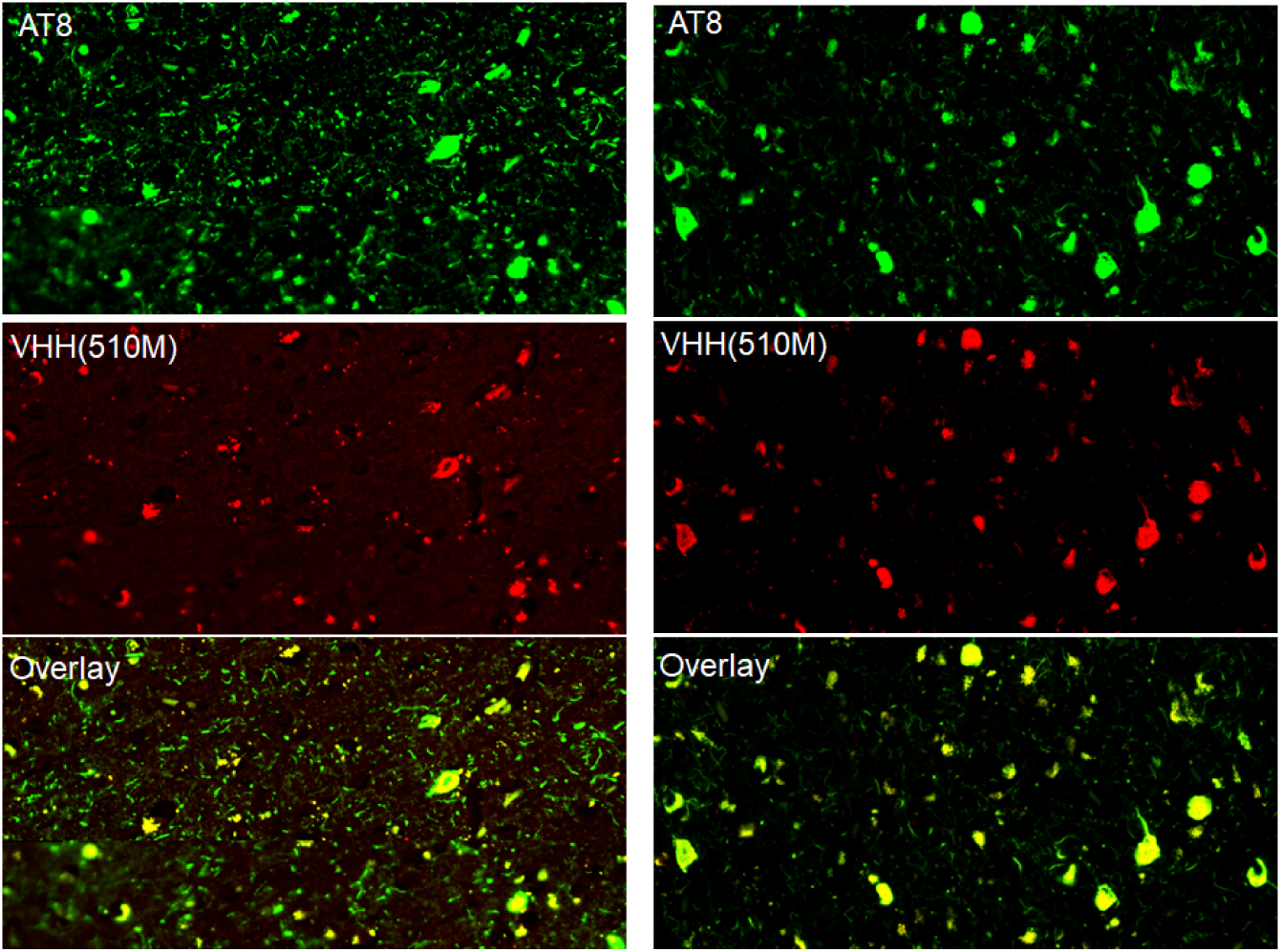
VHH(510M) immunoreactivity in human AD brain tissue. Representative images from tissue microarray sections of fixed frontal cortex of an AD case. Staining with the phospho-tau antibody AT8 (green) and anti-tau VHH(510M) (red) reveals co-localization of tau inclusions in overlapping regions (yellow in overlay). These findings support the selectivity of VHH(510M) for disease-relevant tau aggregates in human AD brains.

## Discussion

Trans-cellular propagation of unique conformations of tau assemblies appears central to Alzheimer’s disease (AD) and other tauopathies (*4*–*6*). Thus, agents that specifically target and neutralize tau seeds, particularly those present in the early stages of the disease, could significantly improve both early detection and therapy. In this study, we focused on generating camelid variable heavy chain (VHH) domains that selectively bind to tau seeds from AD brains. Two major challenges in VHH discovery are the generation of biologically relevant antigens— specifically, the purification of tau seeds from patient brains—and the immunization of llamas or alpacas for VHH library generation, a process that is both time-consuming and labor-intensive. Previous attempts to generate anti-tau VHHs have involved immunizing llamas with small tau peptides, recombinant tau monomer, and heparin-induced tau fibrils and oligomers (*19–23*). To overcome these challenges, we employed a hybrid VHH screening approach to generate seed- selective VHHs.

### Anti-tau VHHs Selectively Target Soluble Tau Seeds from PS19 and Human Tauopathies

Screening directly against soluble lysates that contain native seeds from AD or CBD brains allowed us to identify anti-tau VHH(510M) and VHH(50M)—both displaying high affinity for pathogenic aggregates. Both VHHs selectively bound pathological tau from patient samples vs. recombinant fibrils and had low affinity for tau monomer. Competition experiments indicated that seed binding was unaffected by the presence of excess monomer. The results suggest that these anti-tau VHHs preferentially targeted pathogenic tau seeds. This property is crucial for successful immunotherapy, as the goal is to target pathogenic tau while sparing healthy tau in the cellular environment. This work parallels other efforts to isolate conformation-specific antibodies and nanobodies (*11*, *24*, *24*, *34*, *34–36*), underscoring that subtle structural differences have significant consequences for recognition and therapeutic efficacy.

### VHH(510M) Binds Tau Aggregates in PS19 and AD Brain

Anti-tau VHH bound inclusions present in AD brain tissue. We used the optimized version of anti-tau VHH(510M), which had superior stability and a lower aggregation propensity compared to the VHH(510). Using IP and dot-blot against multiple tauopathy brain homogenates we found that VHH(510M) bound tau seeds from AD and CBD brain but not controls. Finally, immunohistochemistry indicated that the VHH(510M) stained tau inclusions in both PS19 AD brain.

### Potential Applications and Advantages of VHH-based Reagents

Due to their small size, high stability, and ease of genetic manipulation, VHHs hold considerable promise for next-generation immunotherapies and *in vivo* imaging tools (*18*). These nanobodies could be delivered intracerebrally or peripherally, perhaps fused to blood–brain barrier shuttle peptides or packaged within viral vectors, allowing for specific targeting of intracellular tau species. The fact that anti-tau VHH(510M) selectively detects inclusions in both mouse and human AD tissue suggests that it can access pathological assemblies in complex tissue environments. This property may be exploited to visualize early-stage tau lesions via PET imaging or to track the spread of disease-relevant aggregates longitudinally in animal models.

Furthermore, the capacity to distinguish pathogenic tau seeds from monomeric tau might help minimize off-target depletion of physiological tau, which is important for normal neuronal function.

### Challenges and Future Directions

These data illustrate the strong potential of seed-selective VHHs, but several challenges remain. First, it is unclear how effectively VHHs can cross the blood–brain barrier (BBB) *in vivo* without modification. Approaches such as receptor-mediated transcytosis could be employed to improve CNS uptake. Second, while the micromolar affinity for tau monomer is beneficial for seed-specific binding, further engineering may be required to enhance potency or pharmacokinetics if one envisions these VHHs as clinical therapeutics. Protein engineering strategies, such as biparatopic designs or engineered Fc-fusions, might boost avidity and effector function. Third, as tau pathology in human brains is remarkably heterogeneous—encompassing a range of post-translational modifications, structural variants, and strain-specific features—multiple VHH clones with distinct epitopes may prove necessary to achieve robust disease coverage (*20*, *24*, *35*).

### Broadening to other Amyloidogenic Proteins

Many neurodegenerative diseases, including Parkinson’s disease, Huntington’s disease, and amyotrophic lateral sclerosis, are associated with prion-like spread of misfolded proteins such as α-synuclein, huntingtin, and TDP-43. Each of these proteins may exist in structural forms that differ significantly from recombinant assemblies, suggesting that our approach could be generalized to discover VHHs targeting more physiologically relevant conformations. Indeed, early detection and selective neutralization of these misfolded species might represent a powerful strategy for slowing or preventing disease progression.

## Conclusion

We have identified VHHs that preferentially bind pathological tau seeds in AD and CBD, demonstrating utility in cell-based assays, biochemical immunoprecipitations, and histological detection of pathological inclusions. These tools provide new avenues for developing diagnostic probes and potentially disease-modifying therapies that minimize off-target effects on normal tau. Future studies will focus on optimizing VHH engineering for *in vivo* delivery, expanding the breadth of targets, and elucidating how selective interference with pathogenic tau seeds might modify disease trajectory in animal models and ultimately in human clinical settings (*20*, *24*, *35*).

## Materials and Methods

### Library expansion and VHH expression

The "Yeast surface display nanobody library (NbLib)" generated by McMahon et al. (*27*) was procured from Kerafast (Catalog number: EF0014-FP). Upon receipt, the yeast library was thawed and expanded in ‘Yglc4.5 –Trp’ media, as previously described (*27*). Cell viability was assessed and the presence of contamination was ruled out. Finally, we prepared multiple glycerol stocks at >10-fold higher cell viability than the number of clones to ensure no loss in library diversity, and the aliquots were frozen at -80°C for VHH screening.

### VHH expression check

Aliquots of the VHH library were thawed at 30 °C, each containing approximately 5 × 10¹ cells, and were recovered by growing in 1 liter of “–Trp + glucose” media at 30 °C and 220 rpm for 24 hours. The following day, the total number of yeast cells in the overnight culture was counted by measuring the OD_600_ (where OD_600_ of 1 ≈ 1.5 × 10 yeast cells). 1 × 10¹ cells from the “–Trp + glucose” culture was collected and washed once with “–Trp + galactose” media and transferred them into 1 liter of “–Trp + galactose” media. The yeast were grown at 25 °C and 220 rpm for 72 hours, and checked for VHH expression at 24, 48, and 72 hours.

In parallel, 1 × 10¹ cells from the primary culture was transferred into “–Trp + glucose” media as a negative control for VHH expression. At each time point, 1 × 10 cells from both the glucose and galactose cultures was collected, washed twice with 100 µl of selection buffer, and incubated with approximately 0.5 μg of anti-HA antibody labeled with AlexaFluor647 for 30 minutes at 4 °C. The cells were then washed twice with selection buffer to remove unbound antibody and analyzed on a BD LSRFortessa™ Cell Analyzer to determine the percentage of yeast cells expressing nanobodies in both “–Trp + glucose” and “–Trp + galactose” media.

### Preparation of labeled tau for VHH screening

The pet28b plasmid encoding full-length 2N4R tau protein sequence was a kind gift from Dr. David Eisenberg (UCLA). Full-length tau monomer was purified as described previously (*37*), with minor modifications. A pet28a-tau plasmid was transformed into BL21(DE3) competent *E. coli* cells and colonies were screened for protein expression. The colony with the highest protein expression was grown overnight into 50 ml 1×Terrific Broth (TB) media at 37 °C at 220 rpm.

Next day, the primary culture was transferred in 1 liter 1×TB media at 37 °C at 220 rpm and protein expression was induced with 1 mM isopropyl β-D-1-thiogalactopyranoside (IPTG) for 4 h at 37 °C. Cells were harvested by centrifugation at 6000× g for 20 min at 4 °C, resuspended in 50 mM pH 7.5 containing 500 mM NaCl, 1 mM β-mercaptoethanol, 20 mM imidazole, and 1 mM phenylmethylsulfonyl fluoride (PMSF), with cOmplete™, EDTA-free Protease Inhibitor Cocktail, and lysed using GEA PandaPLUS Lab Homogenizer 2000. The cell lysate was centrifuged at 20,000× g for 40 min at 4 °C and the supernatant was filtered and loaded on pre-equilibrated Ni-NTA Agarose beads. The column was washed with lysis buffer and protein was eluted with lysis buffer containing gradient of imidazole from 20 mM to 500 mM. Clean fractions for Ni-NTA purification were concentrated and buffer exchanged into 50 mM MES, 50 mM NaCl, 1 mM β- mercaptoethanol (pH 6.0) by PD-10 column (Cytiva Life Sciences, cat. no 17-0851-01) and loaded onto a5 mL HiTrap SP-HP column (Cytiva Life Sciences, Cat. No. 17115201) for cation exchange purification. Pure fractions from ion-exchange chromatography were pooled, concentrated and injected onto HiLoad 16/600 Superdex 75 pg column (Cytiva Life Sciences, cat. No. 28989333) for size-exclusion chromatography (SEC). The fractions from SEC containing clean protein were pooled and labeled with Alexa-647, Alexa-488, and FITC fluorophore using Alexa Fluor™ 647 NHS Ester (Succinimidyl Ester), Alexa Fluor™ 488 NHS Ester (Succinimidyl Ester), and NHS-Fluorescein (5/6-carboxyfluorescein succinimidyl ester), respectively. For labeling, the purified protein was incubated with a 20-fold molar excess of dye overnight at 4 °C, on a rotatory shaker. The reaction was quenched by adding 0.1 mL of freshly prepared 1.5 M hydroxylamine, pH 8.5 and the free dye was removed by PD10 desalting column. The labeled protein was quantified and stored in -80 °C for further use.

### Magnetic Assisted Cell Sorting (MACS)

Two glycerol stocks of the VHH library containing ∼5x10^9^ cells were thawed and inoculated into 1 liter “–Trp + glucose” media and grown for 24 hours, at 30 °C with shaking at 230 rpm. The OD_600_ was recorded (here, OD_600_ =1 is ∼1x107 cells/ml), and enough yeast for 100-fold library/previous round coverage were inoculated into 1 liter “–Trp + galactose” media and induced for 48 h at 25 °C, 250 rpm. The VHH expression was checked as described above, and yeast was used for MACS. For the first round of MACS, yeast cells were washed twice with selection buffer and the yeast cells were incubated with 500 µl Anti-Cy5/Anti-Alexa Fluor 647 Microbeads (Miltenyi CAT NO. 130-091-395) for 2 h, at 4 °C with constant mixing. The yeast cells were then spun down to remove the unbound magnetic beads and the yeast cells were washed with selection buffer (20 mM HEPES, pH 7.5, 150 mM NaCl, 5 mM maltose, 0.1% BSA) to remove loosely bound magnetic beads. The cells were then passed over a pre-equilibrated LD column (Miltenyi CAT NO. 130-042-901) for negative selection. This step removes all the yeast cells that non-specifically bind to Anti-Cy5/Anti-Alexa Fluor 647 Microbeads. The yeast cells passed through LD column were collected and washed once with selection buffer. The cells were then resuspended and incubated with 500 nM tau2N4R labeled with Alexa647 for 1 hour at 4 °C, with mixing. The yeast cells were then washed twice with selection buffer to remove unbound protein, and the resuspended cells were again incubated with 500 µl Anti- Cy5/Anti-Alexa Fluor 647 MicroBeads (Miltenyi CAT NO. 130-091-395) for 1 h, at 4 °C with constant mixing. This allows formation of yeast-tau-microbead complex that can be selected with MACS. The cells were then washed twice with the selection buffer to remove unbound microbeads and passed throw LS column (Miltenyi CAT NO. 130-042-401) for positive selection. The unbound yeast cells were discarded and the yeast cells collected on LS column were collected and grown in “–Trp + glucose” media for next rounds of screening. We collected a small amount of yeast cells at each step and plated them on “–Trp + glucose agar” and YPD-agar plate to count number of yeast cells after each step, and to confirm no contamination throughout the process. For the second round of MACS, we followed a similar process to the one we used for MACS1. Here, we expanded the yeast cells collected from MACS 1 and expanded them in 1 liter “–Trp + glucose” media for 24 h and then transferred ∼10^9^ yeast cells into 1 liter “–Trp + galactose” media and induced for 48 h at 25 °C, 250 rpm to induce VHH expression. For the second round of MACS, yeast cells were washed twice with selection buffer and the yeast cells were incubated with 500 µl Anti-FITC microbeads (Miltenyi CAT NO. 130- 042-901) for negative selection. The unbound cells were then incubated with 250 nM 2N4R labeled with FITC for 1 hour at 4 °C, while mixing. The yeast cells were then washed and incubated with 500 µl Anti-FITC microbeads (Miltenyi CAT NO. 130-042-901). The yeast cells were then passed through LS column (Miltenyi CAT NO. 130-042-401) for positive selection.

The unbound yeast cells were discarded and the yeast cells collected on LS column were collected and grown in “–Trp + glucose” media for next rounds of screening.

### Fluorescence Activated Cell Sorting (FACS)

Yeast from MACS2 were grown and VHH expression induced as before. After VHH expression was verified, 1x10^7^ yeast were washed in selection buffer and incubated with 100 nM tau 2N4R- AlexaFluor647 for 1 hour, at 4° C with constant mixing. The yeast cells were washed twice with selection buffer to remove unbound tau. The yeast cells bound with tau with then resuspended and incubated with 25 µg of AlexaFluor488-conjugated anti-HA tag antibody (Cell Signaling Technology Cat. No. 2350S). After incubation, yeast cells were washed twice to remove unbound antibody and resuspended in selection buffer and filtered before running on flow sorter. This preparation was diluted 1:5 in selection buffer before being analyzed on a FACSAria SORP 4-laser sorter (BD Biosciences). Yeast cells showing maximal binding to 2N4R-AlexaFluor 647 and anti-HA tag antibody- AlexaFluor488 were collected in “–Trp + glucose” media. Here, we collected ∼0.05% of total cells. Collected yeast cells were revived by growing at 24 hours, at 30 °C with shaking at 230 rpm, and used for the next round of screening.

The cells collected after 1^st^ round of FACS were expanded in 1 liter “–Trp + glucose” media and VHH expression was induced “–Trp + galactose.” The VHH expression was confirmed as described above and cells were used for the 2nd round of FACS. Here, we used 30 nM of 2N4R labeled with FITC was used for screening. The tau binding positive yeast cells were selected and used for the final and 3^rd^ round of FACS. For the final round of FACS we used 10 nM of tau 2N4R labeled with AlexaFluor488 for the screening. After five rounds of screening a total of ∼1000 individual yeast single colonies were collected onto “–Trp + glucose” agar plates and grown for 3-5 days at 30 °C until colonies were visible. These colonies were then screened further to check for their seed binding.

### IC_50_ measurement

The individual yeast colonies were grown in “–Trp + glucose” media and VHH expression was induced “–Trp + galactose” media. A total of 1x10^6^ cells were washed with selection buffer and incubated with various concentrations of fluorescently labeled tau 2N4R for 1 hour, at 4 °C with shaking. Each sample was also incubated with 2 µg fluorophore-conjugated anti-HA antibody for 1 hour, at 4 °C with shaking. The samples were then washed twice with selection buffer and all the samples were then analyzed on LSRFortessa. The percentage of double positive cells with each concentration of labeled tau monomer was plotted and compared for all the VHH clones.

The data was then fitted to a sigmoidal function to derive IC_50_ value.

### VHH sequence verification and cloning in *E. coli* for recombinant protein production

Individual yeast colony were grown in 5 ml “–Trp + glucose” media and DNA was extracted using Zymoprep Yeast Plasmid Miniprep II (cat no D2004). The VHH sequence was then amplified using the following primers: Forward primer: GTTTAACTTTAAGAAGGAGATATACCATGCAGGTGCAGCTGCAGGAAAG Reverse primer: GCCGGATCTCAGTGGTGGTGGTGGTGGTGCTCGAGTTAGCAGCTGCTCACGGTCACCTG.

The PCR product was checked on gel and used for sequencing as well as for cloning in *E. coli* for recombinant protein production. For sequencing, a portion of amplified product was purified with ExoSAP-IT™ (Catalog number 78201.1.ML) and the sequence was determined by Sanger sequencing. For cloning into *E. coli*, the PCR product was cleaned using Zymo Genomic DNA clean and concentrator (cat no D4011). The purified product was then assembled into a pre-digested pET28 vector (digested with NcoI and XhoI enzymes) using NEBuilder® HiFi DNA assembly cloning kit. The assembled product was transformed into NEB® 5-alpha Competent *E. coli* (High Efficiency) cells from NEB. The single colonies were used to purify the plasmid, and the plasmid was sent for sequencing. The VHH sequence was confirmed, and the plasmid was transformed into BL21(DE3) competent cells (prod. no. C2527) for recombinant VHH expression.

### Purification using A3 resin

The pET28a plasmid containing VHH sequence was transformed into BL21(DE3) cells and single colonies were selected and screened for protein expression. The colony having high protein expression was grown overnight in Luria-Bertani broth at 37 °C. The saturated culture was transferred into 1 liter Auto-Induction Medium (cat no GCM17.0500 BOCA Scientific) and grown at 37 °C for 8 hours, followed by overnight at 24 °C, to induce protein expression. The bacterial culture was spun down and cell pellet was resuspended into lysis buffer (1x PBS pH 7.2, 2% glycerol, 2 mM EDTA, protease inhibitor cocktail). The resuspend cells were lysed by GEA PandaPLUS Lab Homogenizer 2000 until clarified. The clarified lysate was spun down to remove cell debris, filtered through 0.45µm filter and purified using Ampsphere A3 resin (JSR Life Sciences). The clarified cell lysate was loaded onto pre-equilibrated Ampsphere A3 resin. The beads were washed with 10 CV (column volumes) of 1x PBS, followed by 10 CV 1M NaCl in PBS, 10 CV 2M NaCl, and finally with 10 CV 5M NaCl to remove non-specifically bound impurities. The nanobodies were then eluted using 2 CV of 100mM Glycine pH 2.5 followed by a second elution using 2 CV 250 mM Glycine pH 2.5. All the elution’s were collected in a tube containing 0.25 CV 1M Tris-HCl pH 8.0 and 0.25 CV 10% glycerol to neutralize the elution’s to avoid VHH precipitation. The VHHs were dialyzed against 1x PBS containing 2% glycerol, filtered using 0.22µm filter and stored at 4 °C for further use.

### Brain lysate preparation

Flash-frozen human or mouse brain was suspended in 1x TBS containing 1x cOmplete protease inhibitor cocktail (Roche) at a final concentration of 10% w/vol. The tissues were homogenized using probe homogenizer with Power Gen 125 tissue homogenizer (Fischer 734 Scientific). The brain lysate was then sonicated for 5 min at amplitude of 65 in a bath-sonicator, with a “30 sec on and 30 sec off” interval to avoid heating of the sample. The sonicated sample was then centrifuged at 20,000 x g, at 4 °C for 20 min, and supernatant was collected in protein low binding tubes. The brain lysate was then quantified using Pierce 660 Assay and used for experiments.

### Immunoprecipitation using Amsphere™ A3 resin

Purified VHHs were concentrated using Amicon ultra centrifugal filter, 3 kDa MWCO (cat. no UFC9003). The VHHs were filtered using Ultrafree-MC 0.22 µm pore size filters (0.5 mL volume, cat. no UFC30GV0S) to remove any aggregated protein. The VHHs were then spun at 20,000 xg for 20 min at 4 °C to remove any residual aggregated VHH and supernatant protein was quantified using DeNovix DS-11 FX+ spectrophotometer. In parallel, clarified the brain lysate was prepared (as described above) and the total protein concentration was quantified using Pierce 660nm protein assay reagent (cat. no 22660). VHHs and lysate were mixed together in a 96-well clear round bottom plate (Corning, cat. no 3788), and incubated overnight at 4 °C, with shaking at 1000 rpm. The following day, 30 µl pre-equilibrated Amsphere A3 beads (prod. no 10000204-330) (bead equilibration process: washing twice with 1x PBS, followed by washing twice with 1x PBST, and twice with TBS + PIC) were added to the “lysate + VHH” reaction mixture. The reaction mixture was incubated for 1 hour, at 4 °C, with shaking at 1000 rpm to allow formation of “bead-VHH-seed complex.” The reaction mixture was then centrifuged at 10,000 xg for 3 min and unbound supernatant was removed and saved for transfection into tau biosensor cells. The beads were then washed four times with TBS+PIC to remove non-specifically bound proteins. Finally, the seeds were eluted by adding 60 µl “Pierce IgG elution buffer” pH 2.5 (cat. no 21004) to the beads. The elution was neutralized by addition of 1/5 part of M Tris-HCl, pH 8.0, and transfected into tau RD(P301S) v2L biosensors, as described previously (*29*). Cells were allowed to grow for 48 hours at 37 °C. After 48 h, the cells were checked for formation of tau puncta in the cells by fluorescence microscopy. The cells were then fixed with 2% PFA, resuspended in 1x PBS and analyzed on LSRFortessa flow cytometer for FRET analysis. The acquired data was processed using FlowJo software, and FRET positive cells were counted and plotted for various samples, as described previously (*38*).

### Immunoprecipitation and monomer competition

For the monomer competition experiment, the brain lysate was supplemented with the purified tau monomer and the immunoprecipitation experiment with Amsphere™ A3 was performed as described above. 10 µg of brain lysate was supplemented with increasing concentrations of purified 2N4R tau monomer and a fixed amount of 10 µg of VHH was then added for IP experiment. Each sample (constant brain lysate concentration with varying tau monomer concentrations), was then incubated overnight at 4 °C with shaking at 1000 rpm. The following day, 30 µl pre-equilibrated Amsphere A3 beads were added to the reaction mixture, and incubated for 1 hour, at 4 °C, with shaking at 1000 rpm. The beads were than washed and the elution was collected by adding 60 µl “Pierce IgG elution buffer” pH 2.5. The elution was neutralized and transfected into tau v2L biosensors, as previously described. The cells were allowed to grow for 48 hours at 37 °C, then fixed with 2% PFA. FRET analysis was performed using an LSRFortessa flow cytometer. The FRET-positive cells at each concentration of tau monomer were compared to assess the effect of the monomer in the brain lysate.

### Dot blot against brain lysates

Clarified brain lysate from two AD cases, one P301S mice and one healthy control were applied onto a polyvinylidene difluoride membrane (Immobilon® -FL PVDF Membrane, Millipore Sigma) using a dot blot apparatus (Bio-Dot Apparatus, BIO-RAD). The membrane was subsequently blocked for 30 min at room temperature in 5% w/v skimmed milk prepared in 1X TBST. The membrane was then incubated overnight at room temperature with 0.001 µg/ml anti-tau VHH(510M) or anti-tau VHH(50M) diluted in 5% w/v skimmed milk prepared in 1X TBST. The membrane was then washed twice with 1X TBST and then incubated with 1:2000 dilution of MonoRab™ Rabbit Anti-Camelid VHH Antibody [HRP], mAb prepared in 1X TBST for 60 min at room temperature. After incubation, the blot was washed twice with 1X TBST for 10 min to wash unbound secondary antibody. Finally, the blot was scanned, and spots were detected using the SuperSignal™ West Femto Maximum Sensitivity Substrate kit, Thermo Scientific. Specific protein signal from the membranes was visualized using a Chemi Doc (Syngene G: BOX Chemi XRQ gel doc system) and images captured with GeneSys Image Capture Software.

### FIDA measurements

Flow Induced Dispersion Analysis (FIDA) measurement for anti-tau VHH(510M) and VHH(50M) were performed on Fida 1 (Fida Biosystems, Denmark) using a dynamic coated (Fida Biosystems, Denmark) 100cm capillary with 75cm length (Fida Biosystems, Denmark). Here, 50 nM of Alexa-647-labeled anti-tau VHHs were incubated with varying concentrations of full-length tau 2N4R. Each sample was then analyzed on the Fida instrument, and the hydrodynamic radius of the complex was measured at each concentration using the 640 nm detector. The parameters of the analysis were:

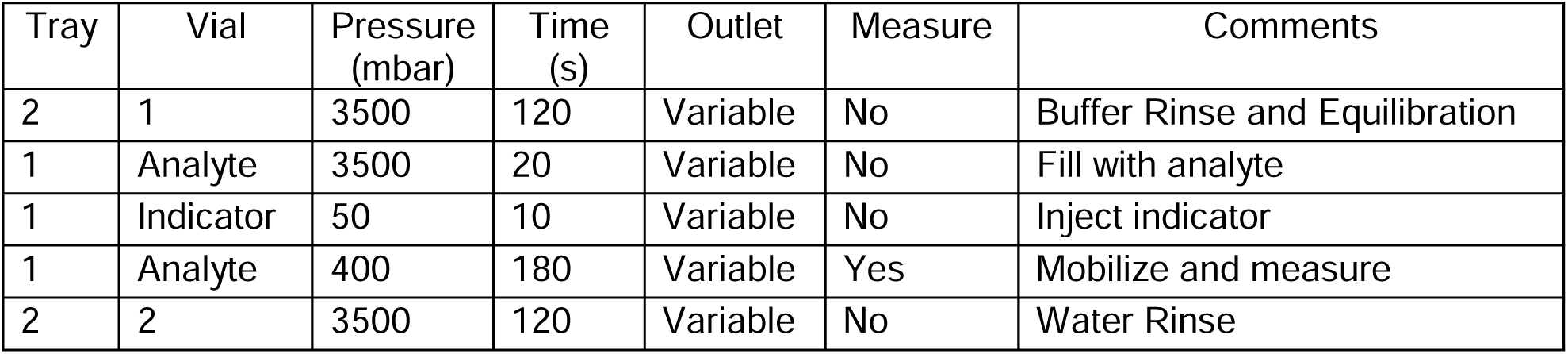

All the trays and capillary were set at 25 °C, the kinetic measurements were also performed at 25 °C.

### CD measurements

Circular dichroism (CD) measurements were performed using a Jasco J-815 spectropolarimeter (Serial No. B064061168) at the UT Southwestern Medical Center in Dallas. Measurements were collected using a 1 mm pathlength quartz cuvette under standard sensitivity settings. A final concentration of 1.0 mg/ml of anti-tau VHH(510M) in 1x PBS containing 2% glycerol was used for all the measurements. The far-UV CD spectra were recorded at 25 °C at a data pitch of 0.1 nm, with a scanning speed of 50 nm/min. The CD thermal melts were recorded from 4 °C to 95 °C by measuring the change in secondary structure at 215 nm (θ_215_), using a CDF-426S temperature control accessory (S/N A00861183). The temperature was controlled with a precision of ±0.10 °C, with a hold time of 5 seconds at each target temperature before data acquisition. The change in θ_215_ was then plotted against temperature and fitted to a sigmoidal function for calculation of mid-point of thermal denaturation (T_m_).

### Preparation of ^15^N tau

BL21(DE3) cells transformed with pET28a plasmid containing tau 2N4R sequence were grown in minimal M9 media for protein production (*39*). The single colony expressing tau protein was grown overnight in M9 media. The following day, the saturated culture was transferred to 1-liter M9 media and protein production was induced with 1mM IPTG for 4 h, at 37 °C. The bacterial culture was centrifuged at 6,000 ×g for 30 minutes, lysed and purified as described above. The pellet was resuspended in 50mM MES pH 6, 10mM EDTA, 10mM DTT, 0.1mM PMSF, with cOmplete™ EDTA-free Protease Inhibitor Cocktail, and lysed using GEA PandaPLUS Lab Homogenizer 2000. The lysate was centrifuged at 15,000× g for 30 minutes, and the supernatant was filtered using a 0.45 μm filter. This clarified lysate was loaded onto a 5 ml HiTrap SP-HP column (Cytiva Life Sciences, cat. No. 17115201), and the protein was purified protein were pooled, concentrated, and injected onto HiLoad 16/600 Superdex 75 pg column (Cytiva Life Sciences, cat. No. 28989333) for SEC. The fractions from SEC containing clean protein were pooled and sent for mass-spectrometric analysis to confirm the degree of isotopic labeling. The ^15^N protein was then stored at -80 °C for NMR experiments.

### NMR measurements

NMR spectra were acquired on an Agilent DD2 spectrometer operating at 800 MHz. ^1^H-^15^N HSQC spectra was performed at 4 °C with samples dissolved in 50 mM sodium phosphate buffer pH6.5 containing 1 mM DTT and 10% D_2_O. A total of 50 µM 2N4R tau protein and 50 µM anti-tau VHH(510M) was used for all the NMR measurements. All data were processed with NMRpipe (*40*) and analyzed with NMRView (*41*).

### Isolation of mouse brain

P301S and WT mice at various ages were anesthetized with isoflurane and perfused with cold PBS. Brains were hemi-dissected. The right hemisphere was frozen in liquid nitrogen and stored at –80 °C for subsequent biochemical assays while the left hemispheres were drop-fixed in phosphate-buffered 4% paraformaldehyde (FD NeuroTechnologies, Colombia, MD, USA) overnight at 4 °C. Left hemispheres were then placed in 10% sucrose in PBS for 24 hours at 4 °C, followed by 24 hours in 20% sucrose in PBS at 4 °C, and finally stored in 30% sucrose in PBS at 4 °C until sectioning.

### Immunohistochemistry of mouse tissue

A sliding-base freezing microtome (Thermo-Fisher Scientific, Waltham, MA, USA) was used to collect 30 μm free-floating coronal sections from fixed mouse brains. The sections were stored in cryoprotectant at 4 °C until immunohistochemistry (IHC) was performed. Slices were washed three times with 1xPBS for 5 minutes, and all subsequent washing steps followed the same procedure. The sections were incubated in 1xPBS containing 0.25% Triton X-100 for 45 minutes at room temperature, for permeabilization. Sections were then incubated with 1 drop of avidin block per 3 ml of IHC blocking buffer, followed by washing. They were subsequently incubated with 1 drop of biotin block per 3 ml in IHC blocking buffer for 1 hour and washed again (Avidin/Biotin Blocking Kit, SP-2001). The sections were incubated with biotinylated AT8 antibody (1:1000, Thermo Scientific) for 4 hours at 25 °C. After washing three times with 1xPBS, the sections were incubated with streptavidin-Alexa Fluor™ 568 conjugate for 1 hour at 25 °C. Following washing, the sections were incubated with a final concentration of 2 ng/µL Alexa Fluor 647-conjugated anti-tau VHH(510M). The next day, the sections were washed, incubated with 1:1000 DAPI for 20 minutes, washed again, and then mounted on charged slides. The slides were allowed to dry overnight before being coverslipped with Aquapolymount. The slides were finally scanned using the Olympus Nanozoomer 2.0-HT (Hamamatsu, Bridgewater, NJ, USA) at the University of Texas Southwestern Medical Center Whole Brain Microscopy Core Facility (RRID:SCR_017949).

### Human tissue staining

Paraffin-embedded human AD and control brain slices were deparaffinated twice with Xylene, each for 5 minutes. Slides were then rehydrated for 2 minutes each in a gradient of 100%, 95%, 70%, and 50% ethanol. Next, the slides containing brain sections were washed under running water for 2 minutes, followed by washing with 1x PBS for 5 minutes, twice. The sections were incubated in 1xPBS containing 0.25% Triton X-100 for 45 minutes at room temperature, for permeabilization. Next, they were blocked in NGS blocking buffer (1XPBST+5% BSA+10% NGS) for 1 hour at room temperature. After blocking the tissues were stained with 1:500 dilution of AT8 in NGS blocking buffer for 4 hours at room temperature. The sections were then washed three times with 1X PBS for 5 minutes and incubated with 1:1000 dilution of streptavidin-Alexa Fluor™ 568 conjugate for 1hr, at room temperature. Finally, the slides were washed three time with 1x PBS and stained with 2 ng/µl of final anti-tau VHH(510M) conjugated with AlexaFluor647 in NGS blocking buffer, overnight at room temperature. The slides were then washed three times with ddH_2_O for 5 minutes each and incubated with 1:1000 DAPI for 20 minutes. The slides were then placed on coverslips with Aquapolymount and allowed to dry. The slide images were acquired on the Olympus Nanozoomer 2.0-HT (Hamamatsu, Bridgewater, NJ, USA) at the University of Texas Southwestern Medical Center Whole Brain Microscopy Core Facility (RRID:SCR_017949).

### Media and buffer used in this study

*Yglc4.5 –Trp (for 1 liter):* Mix 7.6 g of –Trp drop-out media supplement (US Biological D9531) + 6.7 g Yeast Nitrogen Base (Himedia M878) + 10.4 g Sodium Citrate + 7.4 g Citric Acid Monohydrate + 10 mL Pen-Strep (10,000 units/mL stock) + 20 g glucose in sterile Milli-Q water. Once dissolved adjust pH to 4.5 and sterilize the media by filtering through 0.22 µm sterifilter.

*–Trp + glucose media:* Mix 7.6 g of –Trp drop-out media supplement (US Biological D9531) + 6.7 g Yeast Nitrogen Base (Himedia M878) + 6.7 g Yeast Nitrogen Base + 10 mL Pen-Strep (10,000 units/mL stock) + 20 g glucose in sterile Milli-Q. Once dissolved adjust pH to 6.0 and sterilize the media by filtering through 0.22 µm sterifilter.

*–Trp + galactose media:* Mix 7.6 g of –Trp drop-out media supplement (US Biological D9531) + 6.7 g Yeast Nitrogen Base (Himedia M878) + 6.7 g Yeast Nitrogen Base + 10 mL Pen-Strep (10,000 units/mL stock) + 20 g galactose in sterile Milli-Q. Once dissolved adjust pH to 6.0 and sterilize the media by filtering through 0.22 µm sterifilter.

*Selection buffer:* Filter sterilized 20 mM HEPES pH 7.5 buffer with 150 mM sodium chloride, 0.1% (w/v) bovine serum albumin, and 5 mM maltose.

*VHH lysis buffer:* 1x PBS pH 7.2, 2% glycerol, 2 mM EDTA, protease inhibitor cocktail.

*N15 lysis buffer:* 50mM MES pH 6, 10mM EDTA, 10mM DTT, 0.1mM PMSF, protease inhibitor cocktail.

*IHC Blocking buffer:* 5% BSA, 0.25% Triton X-100 (blocking buffer),10% Serum in PBS

## Acknowledgments

We thank Dr. Maikke Ohlson, Dr. Andrea Shiakolas, Nil Saez Calveras, Hesan Jelodari Mamaghani, Dr. Haris Girish, Dr. Sushobhna Batra, Peter Kunach, and Dr. Lukasz A Joachimiak for critical discussions and their suggestions. For cell sorting and flow cytometry instrumentation support, we acknowledge the Moody Foundation Flow Cytometry Facility. We thank Dr. Denise Ramirez and “Whole Brain Microscopy Facility (RRID:SCR_017949)” for microscopy support. NL is supported by the Thomas O. Hicks Scholarship in Medical Research.

## Author Contributions

Conceptualization: AG, JVA, MID Methodology: AG, JVA, MID

Investigation: AG, JVA, RL, DK, VS, YT, KK, SJT, CLW, WPR, JRR, NL

Visualization: AG, JVA, MID Supervision: JVA, MID Writing—original draft: AG, JVA, MID

Writing—review & editing: AG, JVA, RL, DK, VS, YT, KK, SJT, CLW, WPR, JRR, NL

## Data and materials availability

All data needed to evaluate the conclusions in the paper are present in the paper and/or the supplementary materials. Raw data files are available upon request.

