## Supplemental file 1 for "Selective Targeting of Pathogenic Tau Seeds via a Novel VHH"

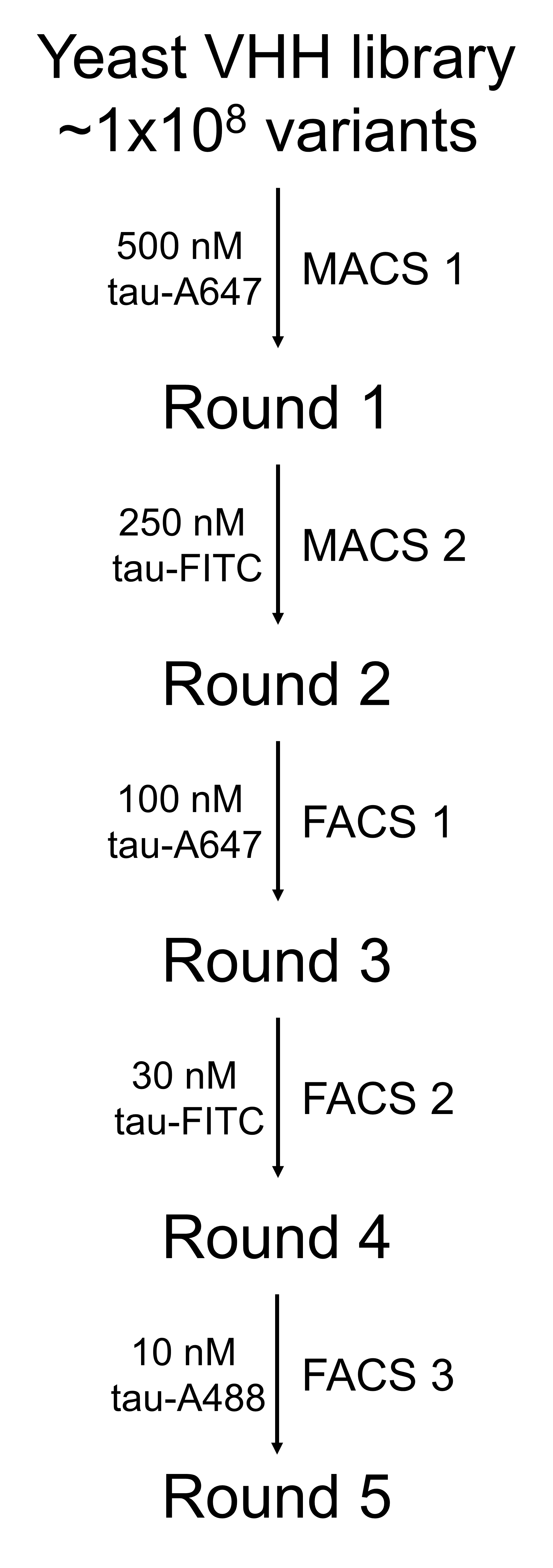

**Figure S1. Initial VHH screening against monomeric tau**

Schematic of the first phase of VHH selection, consisting of five sequential rounds against fill-length tau monomer–two rounds of Magnetic-Activated Cell Sorting (MACS), followed by three rounds of Fluorescence-Activated Cell Sorting (FACS). The tau concentration was progressively lowered at each round to increase selection stringency, and the fluorophore label was changed to avoid screening bias toward a particular dye.

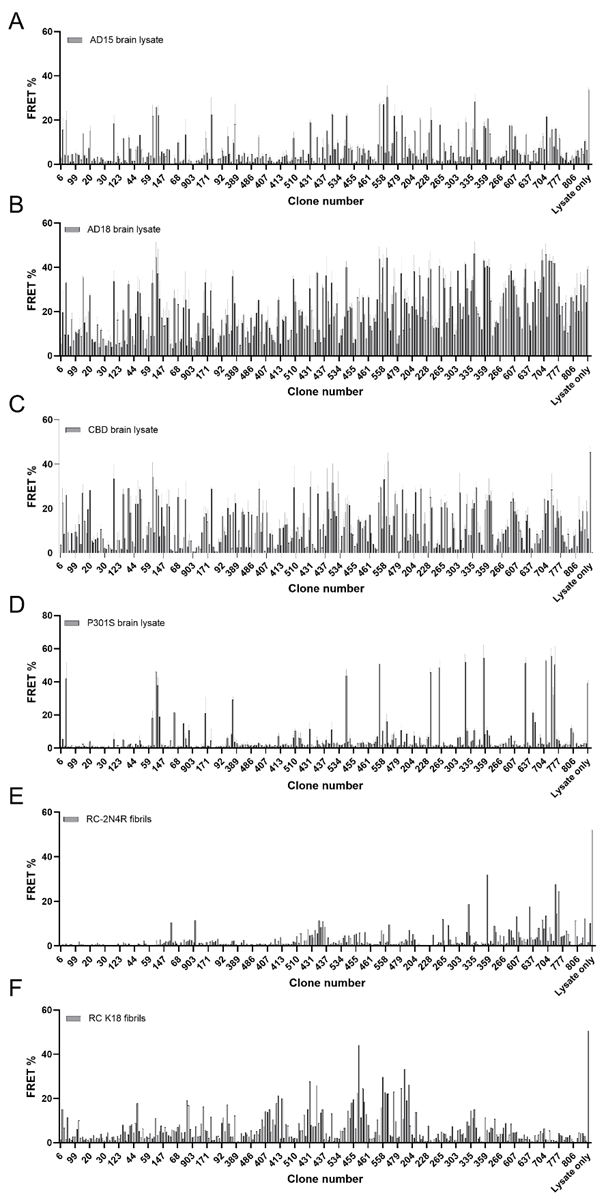

**Figure S2. IP-and-seeding measurement for 300 VHH clones.** Bar plots showing the FRET% readout in V2L tau biosensor cells after immunoprecipitation (IP) with ~300 individual VHH clones. Each VHH was tested against: **(A, B)** Two different AD brain lysates (AD15, AD18), **(C)** CBD brain lysate, **(D)** P301S mouse brain lysate, and **(E, F)** Recombinant heparin-induced 2N4R and K18 tau fibrils. Higher FRET% indicates stronger seeding activity in cells, reflecting greater capacity of a given VHH clone to pull down disease-relevant tau assemblies from each source. “Lysate only” denotes a no-VHH control.

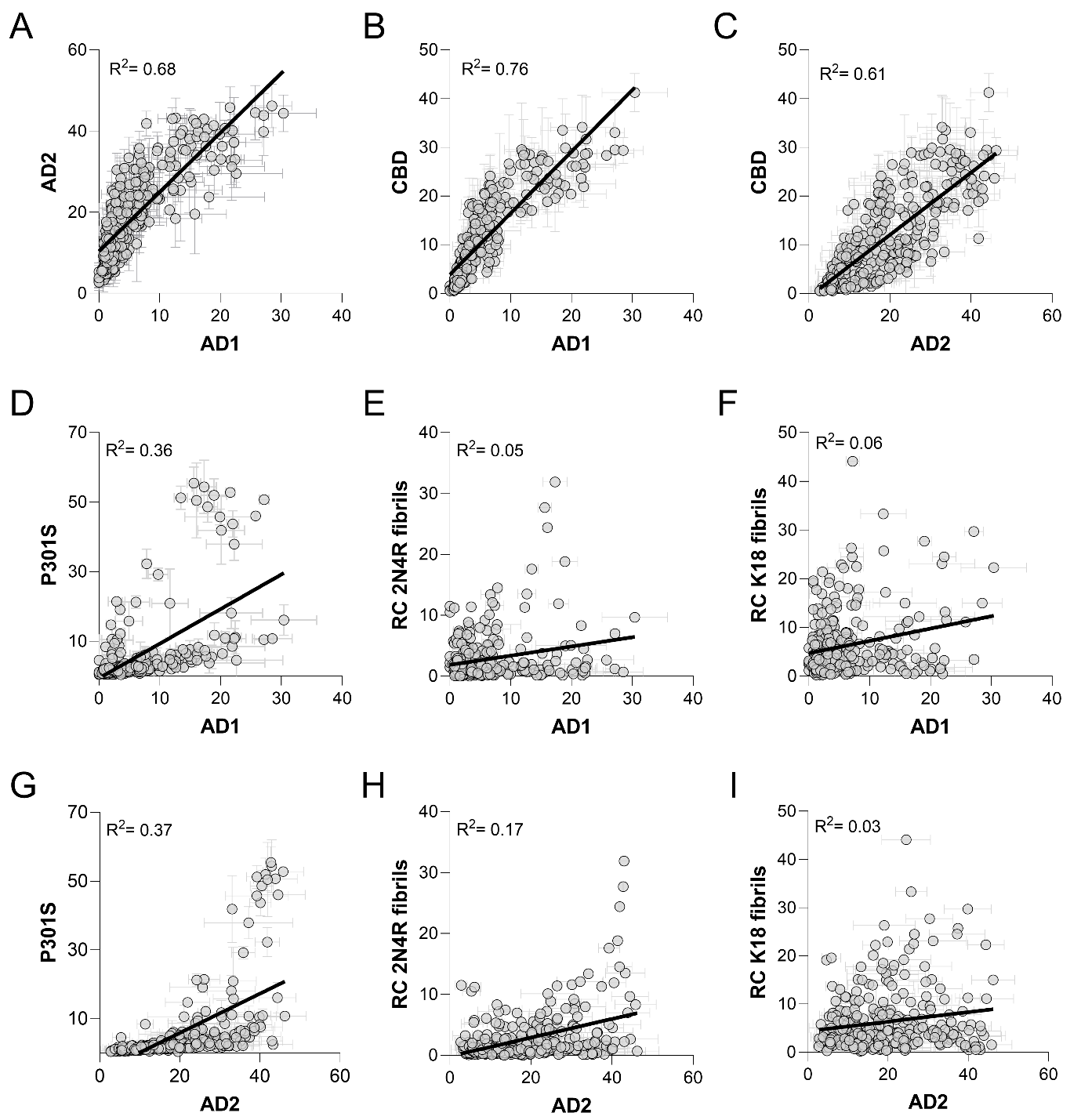

**Figure S3. VHH clones exhibit preferential binding to AD and CBD brain-derived seeds.** Scatter plots compare FRET% values from the same set of 300 top VHH clones tested via IP-and-seeding against multiple tau sources. (**A**) Correlation between two AD cases (AD15 vs. AD18). (**B, C**) Comparisons of AD (AD15 or AD18) and CBD lysates. (**D–F**) Comparisons of AD15 with P301S brain lysate (D) and two types of recombinant fibrils (RC 2N4R, E; K18, F). (**G–I**) Same as (**D–F**), using a different AD brain lysate, AD18. Notably, the strongest correlations occur between the two AD samples and between AD and CBD, whereas correlation with P301S or recombinant fibrils is markedly lower. The top three panels appear in **Figure 2A**.

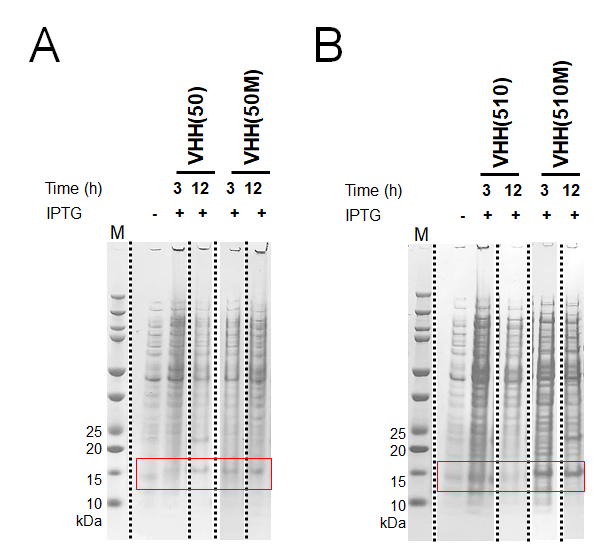

**Figure S4. Recombinant expression of anti-tau VHHs in E. coli.**
(**A**) SDS-PAGE gels showing expression of VHH(50) and VHH(50M). (**B**) Similar analysis for VHH(510) and VHH(510M). In each panel, cultures were induced with 0.25 mM IPTG and harvested after 3 or 12 hours. Cell pellets were lysed in 8 M urea and prepared for SDS-PAGE. The red boxes indicate the expected VHH band around ~15 kDa. M, molecular weight marker; Time (h), induction length in hours.

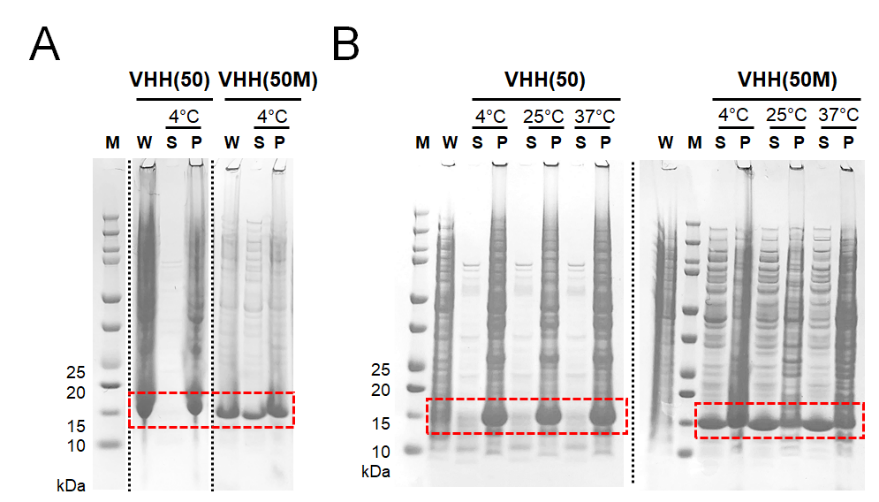

**Figure S5. Temperature-dependent solubility and stability of VHH(50) and VHH(50M).**
SDS-PAGE gels illustrating the impact of temperature on VHH expression and aggregation over time. Cultures were induced for 12 h at 37 °C with 0.25 mM IPTG, followed by lysis in 1×PBS. Both soluble (S) and insoluble (P) fractions were collected and subjected to brief (5 min) or prolonged (48 h) incubation at the indicated temperatures (4 °C, 25 °C, 37 °C) prior to analysis. (**A**) At 5 min post-lysis, wild-type VHH(50) was largely found in the pellet (P), whereas VHH(50M) showed partial solubility. (**B**) After 48 h, the soluble fraction of VHH(50M) remained stable at all temperatures, whereas wild-type VHH(50) exhibited extensive aggregation. Red boxes highlight the expected ~15 kDa VHH band. M: molecular weight marker; W: whole cell lysate (solubilized in 8 M urea).

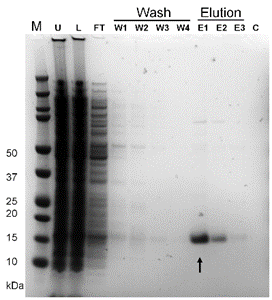

**Figure S6. Purification of anti-tau VHH(510M) using Amsphere™ A3 resin.**
SDS-PAGE gel illustrating each purification step. M, molecular weight marker; U, whole cell lysate in 8 M urea; L, clarified cell lysate loaded onto the A3 column; FT, flow-through containing unbound material; W1–W4, successive wash steps with 1xPBS containing 500 mM, 1 M, 2 M, and 5 M NaCl, respectively; E1–E3, elutions collected with 200 mM glycine (pH 2.5); C, wash with 1 M NaOH. The purified VHH(510M) (~15 kDa, arrow) elutes in E1 and E2. Pooled fractions were subsequently subjected to size-exclusion chromatography (SEC) to remove high-molecular-weight impurities, and the final purified protein was used in downstream experiments.

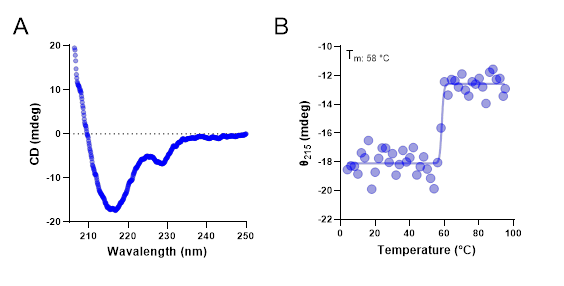

**Figure S7. Confirmation of VHH(510M) folding and stability.** (**A**) Far-UV circular dichroism (CD) spectrum of purified VHH(510M), showing a characteristic β-sheet minimum around 215 nm. (**B**) Thermal denaturation profile (4 °C to 95 °C), monitored by the change in ellipticity at 215 nm (θ_215_). VHH(510M) remains stably folded up to approximately 58 °C, beyond which it transitions to an unfolded state. Data in B are also shown in **Figure 6A** of the main text.

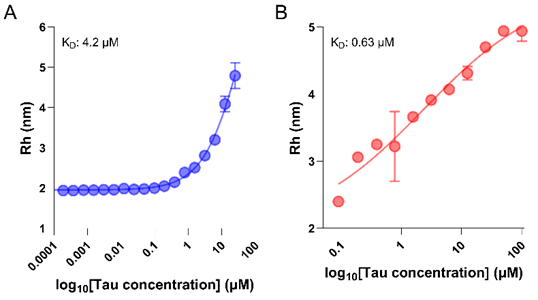

**Figure S8. In-solution affinity measurements of VHH(510M) and VHH(50M).**
(**A**) Binding of VHH(510M) to tau monomer measured using Flow-Induced Dispersion Analysis (FIDA). The hydrodynamic radius (R_h_) of the VHH–tau complex is plotted against various tau concentrations, yielding a K_D_ of ~4.2 µM upon fitting to a 1:1 binding model. (**B**) Similar analysis for VHH(50M), yielding a K_D_ of ~0.63 µM. Data in A are also shown in **Figure 6B** of the main text.

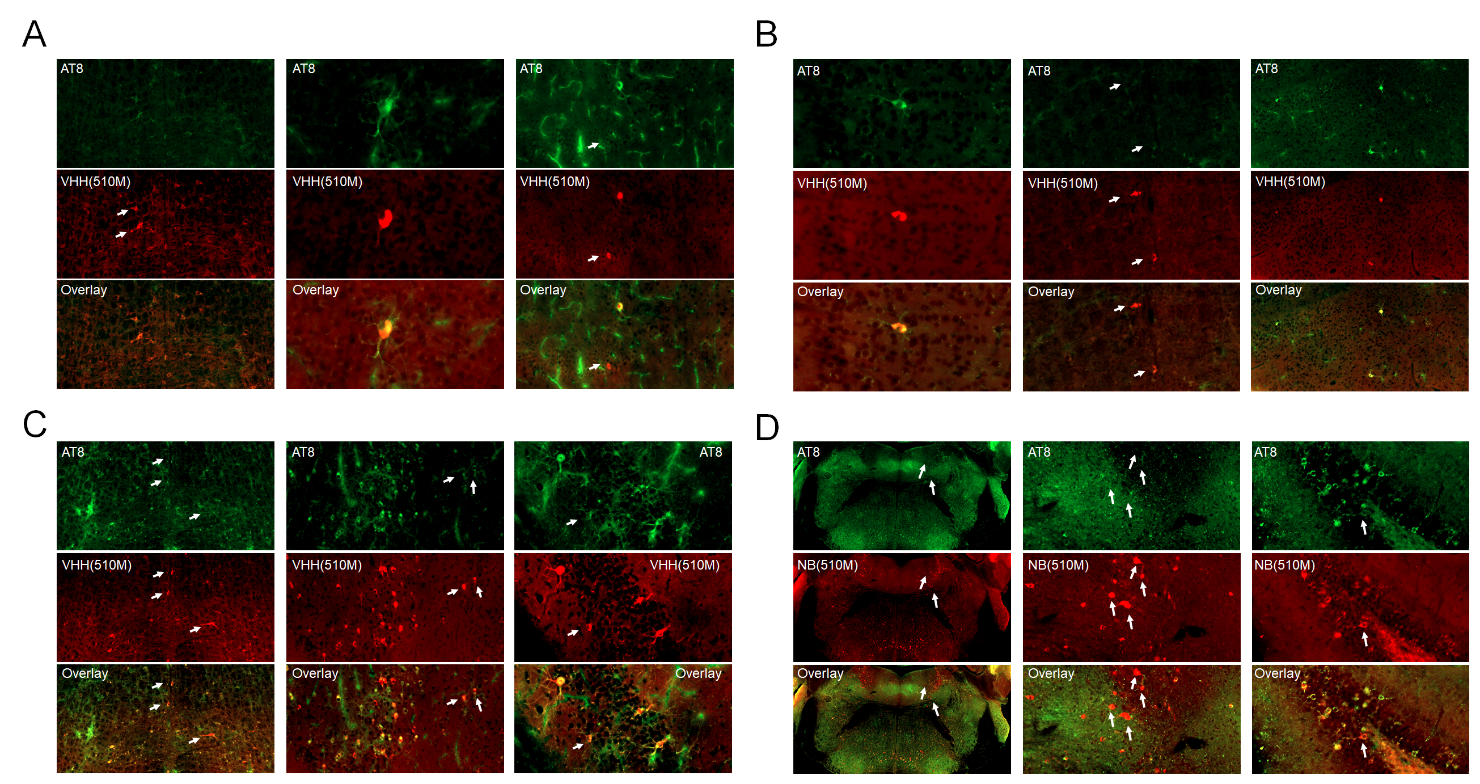

**Figure S9. Immunofluorescent staining of P301S mouse brain sections with VHH(510M).**
Representative brain sections from P301S tauopathy mice at (A) 3, (B) 6, (C) 9, and (D) 12 months of age stained with phospho-tau antibody AT8 (green) and anti-tau VHH(510M) (red). The white arrows denote inclusions recognized by VHH(510M) that are not visibly stained by AT8, suggesting that VHH(510M) detects additional tau aggregates beyond phosphorylated species. Data shown in the last column of panels A, C and D, and the last column of panel B are also shown in **Figure 7**.
